# Quantitative Model of the Ocular Immune Response during Seasonal Allergic Conjunctivitis

**DOI:** 10.64898/2026.09.21.753366

**Authors:** Narshini D. Gunputh, Kara L. Maki, Lucia Carichino, Collynn F. Woeller

## Abstract

Allergic conjunctivitis is an inflammation of the conjunctiva caused by allergen; it is common disorder affecting up to 40% of the population. In this work, we study seasonal allergic conjunctivitis (SAC), also called “hay fever eyes”, which is caused by exposure to airborne pollens. We develop a mathematical model quantifying the ocular immune system response to the allergens. First, we present a simplified qualitative description of the immunopathogenesis of SAC. Then, we express each chosen immunopathological mechanism mathematically to construct a system of thirty-one ordinary differential equations. We compare summary statistics of the predicted observable immune signals to experimental measurements and find our model captures key qualitative features of SAC progression. We then compare our predicted time series of histamine concentration to symptom scores and find a strong correlation suggesting the model predicts relevant clinically trends. Next, we calibrate the model through multi-step process. We find the most influential parameters are the production and depletion rates of IL-4, and the production rates of IL-5 and IL-8. These cytokines are targeted in treatments for asthma, atopic dermatitis, and severe eosinophilic associated disorder and suggest potential therapeutic targets for SAC. Our calibrated model mimics most of the summary statistics of the experimentally observable immune signals with discrepancies for IL-5 and IL-13 indicating that additional immunopathological mechanisms could be important.

## 1 Introduction

Seasonal allergic conjunctivitis (SAC) is one of the most common ocular inflammatory diseases worldwide; it is characterized by itching, hyperemia, tearing, and discomfort driven by a dysregulated immune response at the ocular surface. Unlike many systemic allergic conditions, inflammation in the conjunctiva must be tightly controlled to preserve tissue integrity and visual function [1, 2]. This balance is achieved through a complex interplay between epithelial cells, innate immune populations, adaptive T-helper responses, and a network of pro- and anti-inflammatory cytokines operating within the tear and conjunctival compartments [1, 3].

Immunologically, SAC is dominated by a Th2-mediated response involving allergen-driven activation of antigen-presenting cells, differentiation of naive T-helper cells into allergen-specific Th2 cells, IgE class switching in B cells, mast cell degranulation, and recruitment of eosinophils [1]. These processes are further modulated by regulatory cytokines such as IL-10 and TGF-*β*, which act to dampen excessive inflammation without fully suppressing immune activity [4–6]. Experimental and clinical studies have measured elevated levels of IL-4, IL-5, IL-6, IL-8, IL-13, TNF-*α*, histamine, and IgE in the tears of patients with allergic conjunctivitis [7]. Unfortunately, the interactions between immune cells and inflammatory mediators governing the progression and regulation of conjunctival inflammation remain poorly understood at a systems level [8].

Mathematical modeling provides a framework for integrating these interacting pathways and for interrogating the relative contributions of immune cells and signals to disease dynamics. Prior ophthalmological mathematical models have largely focused on ocular drug delivery, pharmacokinetics, and transport phenomena, including compartmental and physiologically-based pharmacokinetic models describing drug distribution across ocular tissues [9, 10]. While such models have been instrumental in guiding dose selection and therapeutic design, they do not explicitly resolve the immune-mediated mechanisms underlying allergic inflammation at the ocular surface [11].

In contrast, mechanistic models of immune dynamics have been widely applied to systemic allergic diseases, asthma, and autoimmune disorders [12]. Clinical and epidemiological studies suggest that allergic conjunctivitis shares key immunological and physiological features with respiratory allergic diseases [13]. In particular, Michailopoulos et al. reported a strong association between allergic conjunctivitis and asthma, supporting the concept of a shared underlying allergic immune response across ocular and airway tissues [13]. Both tissues possess mucosal surfaces rich in goblet cells and IgE-mediated immune components, facilitating allergen penetration and triggering similar Th2-driven inflammatory cascades. These shared structural and immunological features motivate the use of established asthma and respiratory allergy models to inform mechanistic representations of immune activation in SAC [13].

In this work, we develop a mechanistic model describing the immune response in the conjunctiva during SAC. The objectives of this modeling effort are threefold. First, we aim to construct a biologically grounded mathematical model capable of reproducing key features of conjunctival immune activation in response to allergen exposure. Second, we seek to characterize the relative influence of immune pathways and parameters on inflammatory outcomes with the aim of identifying influential parameters. Third, we calibrate the model parameters to evaluate which processes can be reliably inferred from observable quantities and to guide model reduction and interpretation. By providing a quantitative framework for immune regulation at the ocular surface, this work contributes to the growing body of mathematical immunology models tailored to tissue-specific inflammation and used to identify potential therapeutic targets.

In what follows, we present the mathematical model in Section 2 followed by a description of the methods used to simulate and calibration the model in Section 3. We then present our results in Section 4, and end with our discussion and conclusions in Section 5.

## 2 Mathematical Model

We develop a mathematical model to describe the immunopathogenesis of SAC. The model captures immune activity localized to the ocular surface, and does not include systemic circulation dynamics. It describes the dynamics of key immune cell populations in the conjunctival microenvironment along with the immune signaling network following repeated exposure of allergens in the tears over an allergy season. In what follows, we first introduce our simplified qualitative description of the ocular surface’s adapted immune response to seasonal allergies. Then, we present our quantitative description of immune respond followed by our complete mathematical model.

### 2.1 Qualitative Description of the Immune Response in SAC

We model the abnormal immune response causing acute inflammation in SAC. We consider the dynamics of cell populations in the epithelial layer, and the subepithelial layer containing the fibrovascular connective tissue and immune cells of the conjunctiva [14]. More specifically, in the subepithelial layer, we consider the antigen-presenting cell populations (dendritic cells and macrophages), the adaptive immune cell populations (T-helper 2 or Th2 cells, B cells, and plasma cells), and the effector cell populations (mast cells and eosinophils). Additionally, we model the dynamics environmental and immune signals within the tissue. A complete of model dependent variables are given in Table 1.

**Table 1.**
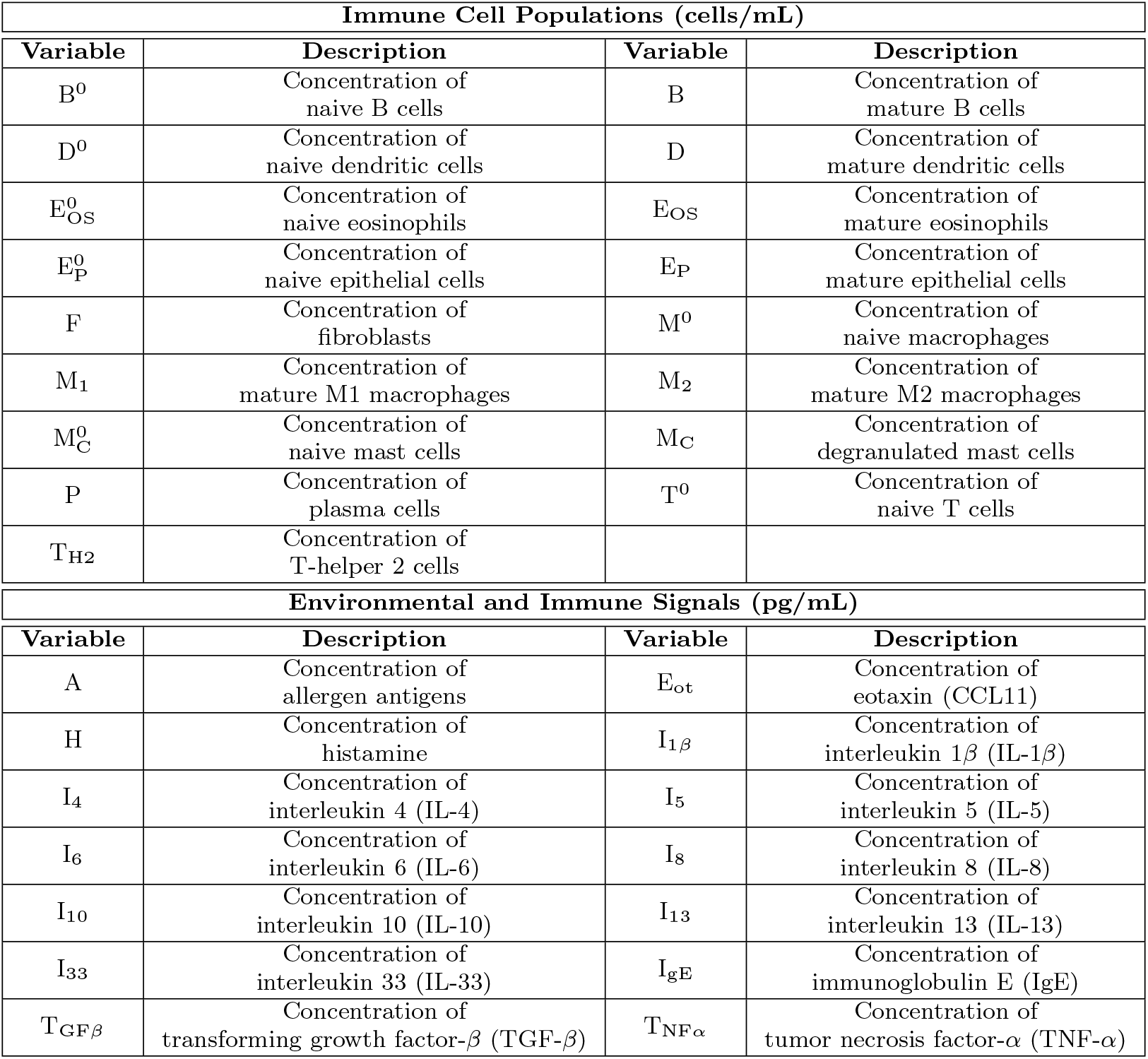
Description of the model dependent variables. Immune cell populations are listed first in alphabetical order followed by the environmental and immune signals.

| Immune Cell Populations (cells/mL) |  |  |  |
| --- | --- | --- | --- |
| Variable | Description | Variable | Description |
| $B^0$ | Concentration of naive B cells | B | Concentration of mature B cells |
| $D^0$ | Concentration of naive dendritic cells | D | Concentration of mature dendritic cells |
| $E_{OS}^0$ | Concentration of naive eosinophils | $E_{OS}$ | Concentration of mature eosinophils |
| $E_P^0$ | Concentration of naive epithelial cells | $E_P$ | Concentration of mature epithelial cells |
| F | Concentration of fibroblasts | $M^0$ | Concentration of naive macrophages |
| $M_1$ | Concentration of mature M1 macrophages | $M_2$ | Concentration of mature M2 macrophages |
| $M_C^0$ | Concentration of naive mast cells | $M_C$ | Concentration of degranulated mast cells |
| P | Concentration of plasma cells | $T^0$ | Concentration of naive T cells |
| $T_{H2}$ | Concentration of T-helper 2 cells | | |
| Environmental and Immune Signals (pg/mL) |  |  |  |
| Variable | Description | Variable | Description |
| A | Concentration of allergen antigens | $E_{ot}$ | Concentration of eotaxin (CCL11) |
| H | Concentration of histamine | $I_{1\beta}$ | Concentration of interleukin $1\beta$ (IL-$1\beta$) |
| $I_4$ | Concentration of interleukin 4 (IL-4) | $I_5$ | Concentration of interleukin 5 (IL-5) |
| $I_6$ | Concentration of interleukin 6 (IL-6) | $I_8$ | Concentration of interleukin 8 (IL-8) |
| $I_{10}$ | Concentration of interleukin 10 (IL-10) | $I_{13}$ | Concentration of interleukin 13 (IL-13) |
| $I_{33}$ | Concentration of interleukin 33 (IL-33) | $I_{gE}$ | Concentration of immunoglobulin E (IgE) |
| $T_{GF\beta}$ | Concentration of transforming growth factor-β (TGF-β) | $T_{NF\alpha}$ | Concentration of tumor necrosis factor-α (TNF-α) |

The immunopathogenesis of SAC comprises three phases referred to as (i) sensitisation, (ii) early, and (iii) late phases [14]. Our mathematical model captures aspects of each of these phases. Figure 1 shows a schematic summarizing all the immunopathological mechanisms or immunopathways described in the model. Our model includes the key features of an overactive immune response to environmental allergens according to established experimental and clinical literature (reference in Table 2). We note that not all immunopathways are included in the model.

**Table 2.** Description of the different interactions of the immune cells and inflammatory mediators in the model.

| Pathway Number | Symbolic Representation | Mechanism Type | Justification |
| --- | --- | --- | --- |
| 1 | $\text{Ep}^0 \rightarrow \text{Ep}$ (A) | Activated change-of-state | Allergens activate the conjunctival epithelial cells as the first cellular target. Epithelial cells are activated into mature epithelial cells and releases pro-inflammatory cytokines [18]. Epithelial cells initiate and actively participate in inflammation [19]. |
| 2 | $\text{D}^0 \rightarrow \text{D}$ (A) | Activated change-of-state | Allergens active naive dendritic cells into mature dendritic cells [1, 3]. |
| 3 | $\text{M}^0 \rightarrow \text{M}_1$ (A) | Activated change-of-state | Allergens activates the naive macrophages driving their polarization to M1 macrophages [3, 20]. |
| 4 | $\text{B}^0 \rightarrow \text{B}$ (A) | Activated change-of-state | Allergens activate the naive B cells into mature B cells [17, 21]. |
| 5 | $\text{B} \rightarrow \text{P}$ (A) | Activated change-of-state | Allergens drive the differentiation of activated B cells into IgE-secreting plasma cells [17]. |
| 6 | $\text{D} \uparrow \text{I}_6$ | Production | Mature dendritic cells secrete IL-6 [22]. |
| 7 | $\text{M}_1 \uparrow \text{I}_6$ | Production | M1 macrophages secrete IL-6 (primary source in allergic inflammation) [22]. |
| 8 | $\text{Ep} \uparrow \text{I}_{33}$ | Production | Mature epithelial cells release IL-33 [3]. |
| 9 | $\text{Ep} \uparrow \text{I}_8$ | Production | Mature epithelial cells produce IL-8 [3]. |
| 10 | $\text{T}_\text{H}^0 \rightarrow \text{T}_\text{H2}$ ($\text{I}_6$) | Activated change-of-state | IL-6 activates naive T cells into Th2 cells [23, 24]. |
| 11 | $\text{T}_\text{H}^0 \rightarrow \text{T}_\text{H2}$ ($\text{I}_{33}$) | Activated change-of-state | IL-33 enhances the activation of Th2 cells via the receptor ST2 [3, 12]. |
| 12 | $\text{T}_\text{H}^0 \rightarrow \text{T}_\text{H2}$ ($\text{I}_{1\beta}$) | Activated change-of-state | IL-1β enhances the activation of Th2 cells [25]. |
| 13 | $\text{E}_\text{OS}^0 \rightarrow \text{E}_\text{OS}$ ($\text{I}_{33}$) | Activated change-of-state | IL-33 activates the eosinophils cells [26]. |
| 14 | $\text{T}_\text{H}^0 \rightarrow \text{T}_\text{H2}$ ($\text{I}_4$) | Activated change-of-state | IL-4 enhances the activation of Th2 cells through a feedback loop [1]. |
| 15 | $\text{T}_\text{H}^0 \not\rightarrow \text{T}_\text{H2}$ ($\text{T}_{\text{GF}\beta}$) | Inhibited change-of-state | TGF-β inhibits the activation of Th2 cells [27]. |
| 16 | $\text{T}_\text{H2} \uparrow \text{I}_{10}$ | Production | Th2 cells produce IL-10 [1, 3]. |
| 17 | $\text{M}_2 \uparrow \text{I}_{10}$ | Production | M2 macrophages upregulate the production of IL-10 [28, 29]. |
| 18 | $\text{T}_\text{H2} \uparrow \text{I}_5$ | Production | IL-5 is produced by Th2 cells [1, 3]. |
| 19 | $\text{T}_\text{H2} \uparrow \text{I}_{13}$ | Production | IL-13 is produced by Th2 cells [1, 3]. |
| 20 | $\text{T}_\text{H2} \uparrow \text{I}_4$ ($\text{I}_{13}$) | Activated production | IL-4 production by Th2 cells is enhanced by IL-13. IL-13 acts through the shared IL-4Rα/IL-13Rα1 receptor on Th2 cells that engages STAT6 and drives GATA3 expression which further enhances the production of IL-4 [1, 3, 30, 31]. |
| 21 | $\text{T}_\text{H2} \not\uparrow \text{I}_4$ ($\text{I}_{10}$) | Inhibited production | IL-10 directly inhibits IL-4 production by Th2 cells via IL-10 receptor (IL-10R). IL-10 acting through IL-10R/STAT3 on Th2 cells limits their antigen-specific activation and directly suppresses secretion of IL-4 [1, 32, 33]. |
| 22 | $\text{E}_\text{OS} \uparrow \text{I}_4$ | Production | IL-4 is produced by mature eosinophil cells [3]. IL-4 is pre-formed in eosinophils and rapidly released during activation which amplifies chronic allergic response [34]. |
| 23 | $M_C \uparrow I_4$ | Production | IgE-activated mast cells produce IL-4 upon degranulation [1, 3]. |
| 24 | $F \uparrow E_{ot} (I_4)$ | Activated production | Fibroblasts produce eotaxin/CCL11 in response to TNF-α signaling through the TNFR1 [3, 35]. |
| 25 | $F \uparrow I_8 (I_4)$ | Activated production | IL-4 upregulates the production of IL-8 from fibroblasts [36]. |
| 26 | $F \uparrow I_8 (T_{NF\alpha})$ | Activated production | TNF-α induces the release of IL-8 from fibroblasts [3]. |
| 27 | $E_{OS}^0 \rightarrow E_{OS} (E_{ot})$ | Activated change-of-state | Eotaxins, through the CCR3 receptor, activate eosinophils cells [37]. |
| 28 | $E_{OS}^0 \rightarrow E_{OS} (I_5)$ | Activated change-of-state | IL-5 enhances the differentiation of eosinophil cells through the IL-5Rα receptor [38]. |
| 29 | $M_1 \uparrow I_{1\beta}$ | Production | Upon polarization, M1 macrophages produces IL-1β [3]. |
| 30 | $B^0 \rightarrow B (I_4)$ | Activated change-of-state | IL-4 activates the proliferation of B-cells [3]. |
| 31 | $M_C^0 \rightarrow M_C (I_{10})$ | Inhibited change-of-state | IL-10 stabilizes mast cells by decreasing FcεRI expression limiting mediator release, and suppressing degranulation [39, 40]. |
| 32 | $M_C^0 \rightarrow M_C (I_{gE})$ | Activated change-of-state | IgE activates the granulation of mast cells through the crosslinking of the antibody IgE to the mast-cell receptor FcεRI [1, 3]. |
| 33 | $M_1 \uparrow T_{NF\alpha} (I_{10})$ | Inhibited production | IL-10 reduces TNF-α release from IgE-activated mast cells [41]. |
| 34 | $M_C \uparrow T_{NF\alpha}$ | Production | Degranulated mast cells produces TNF-α [3]. |
| 35 | $M_1 \uparrow T_{NF\alpha}$ | Production | M1 macrophages release TNF-α [42, 43]. |
| 36 | $M_C \uparrow H$ | Production | Histamine is produced by degranulated mast cells [1, 3]. |
| 37 | $B \rightarrow P (I_4)$ | Activated change-of-state | IL-4 drives the differentiation of B cells into IgE-secreting plasma cells [1]. |
| 38 | $P \uparrow I_{gE}$ | Production | Plasma cells produce IgE [1]. |
| 39 | $M_2 \uparrow T_{GF\beta}$ | Production | M2 macrophages produces TGF-β [44, 45]. |
| 40 | $M^0 \rightarrow M_2 (I_4)$ | Activated change-of-state | IL-4 activates the naive macrophage cells into M2 macrophages [46]. |

**Fig 1.**
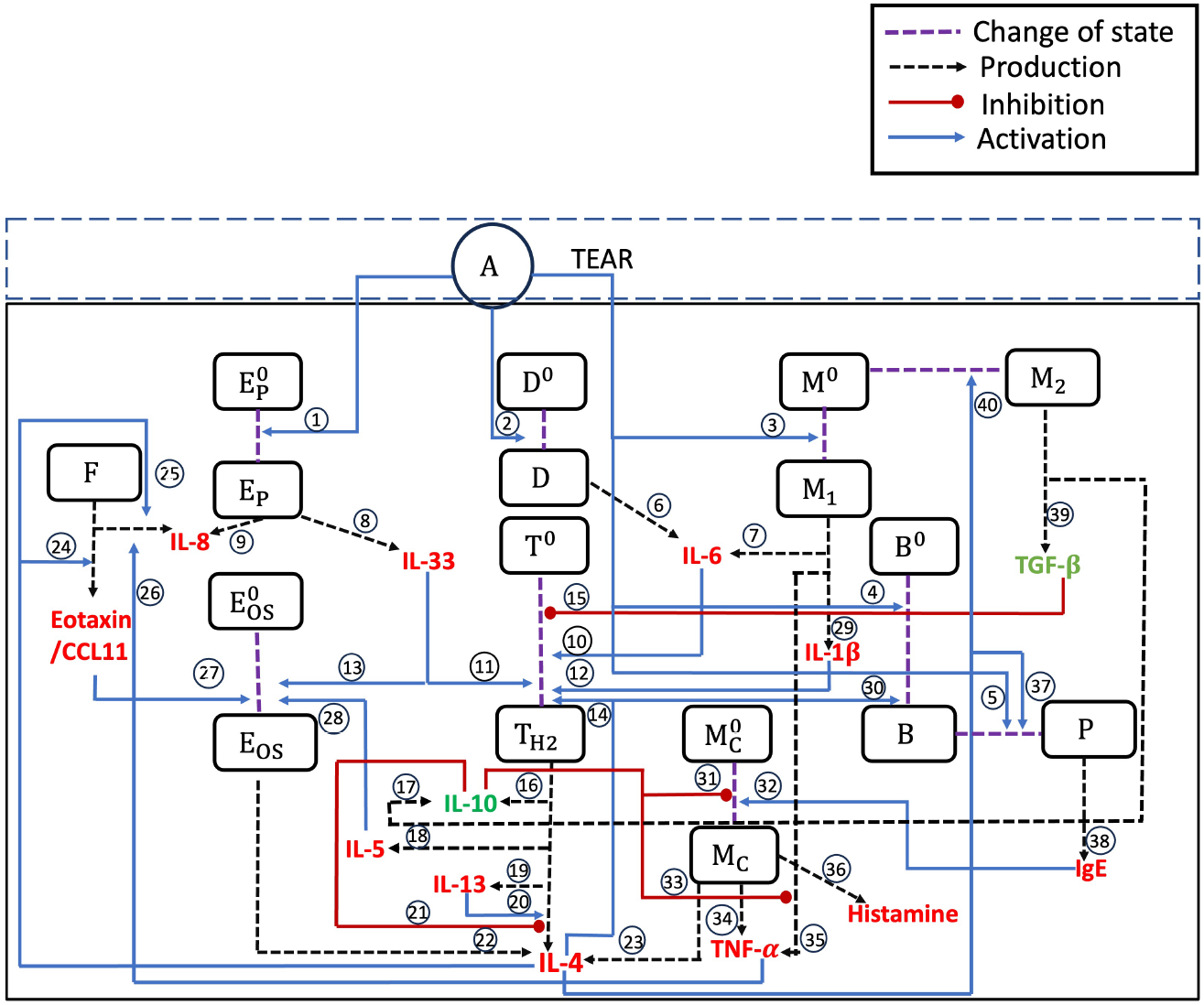
Schematic representing the dynamics of population interactions. The dotted lines represent a change in state from an inactive to an activated state (purple) or a production of an inflammatory mediator or another cell (black arrow). The solid line either represent a boost in activation (blue arrow) or an inhibition (red circle). The pathways describing the interactions are represented by numbers 1-40.

The type of immunopathological mechanisms considered include the (i) activated cell change-of-state, (ii) inhibited cell change-of-state, (iii) activated immune signal production, (iv) immune signal production, and (v) inhibited immune signal production. Each mechanism or pathway is numbered and will be referred to accordingly below. In what follows, we describe the immunopathways included on our model. The biological justification for each pathway is given in Table 2.

To model the sensitisation phase, we assume allergen antigens present in the tears interact with the naive epithelial cell population (pathway 1) or penetrate the conjunctival epithelial barrier entering the subepithelial layer interacting with the following naive immune cell populations: dendritic cells, macrophages, and B cells (pathways 2-4, respectively). The penetration of the conjunctival epithelial barrier is due to the presence of globlet cells and permeable mucosal barrier [15, 16]. Consequently, these naive cell populations are activated into mature cell populations that can produce cytokines (protein signals) or activate other immune cells [1, 3, 17]. If allergen antigens are present, mature B cells differentiates into plasma cells (pathway 5).

Most critically, naive T cells are activated into Th2 cells (pathways 10, 11, and 12) via cytokine interleukin 6 (IL-6) produced by activated dendritic cells and M1 macrophages (pathways 6 and 7); IL-1*β* produced by activated M1 macrophages (pathway 29); and IL-33 produced by activated epithelial cells (pathway 8). Active Th2 cells produce cytokines IL-4 (pathway 21), IL-5 (pathway 18), IL-10 (pathway 16), and IL-13 (pathway 19).

We assume IL-4 is central to the immune regulation [1]. IL-4 enhances the activation of Th2 cells (pathway 14), of M2 macrophages (pathway 40), and of mature B cells (pathway 30). IL-4 also proliferates plasma cells (pathway 37). The plasma cells produced immunoglobulin E (IgE) (pathway 38). Characteristics of the early phase include the release of histamine. In our model, the IgE triggers mast cell deganulation (pathway 32). Consequently, the degranulated mast cells produce histamine (pathway 36), IL-4 (pathway 23) and TNF-*α* (pathway 34). TNF-*α* enhances the production of IL-8 by the fibroblasts (pathway 26). Similarly, IL-4 enhances the production of exotaxin/CCL11 (pathway 24) by the fibroblasts. The late phase of SAC is captured via the recruitment of eosinophils via IL-5 (pathway 28), IL-33 (pathway 13), and exotaxin/CCL11 (pathway 27). Eosinophils produce more IL-4 (pathway 22).

We also model complex interplay of immune signal network. For example, IL-10 inhibits the production of IL-4 by Th2 cells (pathway 21); the production of TNF-*α* by M1 macrophages (pathway 33); and the deganulation of mast cells (pathway 31). TGF-*β* inhibits the activation of naive T cells to Th2 cells (pathway 15).

The biological justification for each pathway is summarized in Table 2. The symbolic representation is such that *V*_1_ *→ V*_2_(*V*_3_) indicates that variable *V*_1_ changes state to variable *V*_2_ when activated by variable *V*_3_. On the other hand, 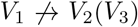 indicates that variable *V*_3_ inhibits variable *V*_1_ changing state to variable *V*_2_. Then *V*_1_ *↑ V*_2_ indicates that variable *V*_1_ produces variable *V*_2_. If *V*_1_ *↑ V*_2_(*V*_3_), then variable *V*_3_ actives the production of variable *V*_2_ by variable *V*_1_. Finally, if 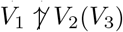, then variable *V*_3_ inhibits the production of variable *V*_2_ by variable *V*_1_.

### 2.2 Quantitative Description of the Immune Response in SAC

We assume the allergens, immune cell populations, and immune signals are spatial heterogeneity. This assumption is consistent with prior immune modeling studies in which spatial resolution is secondary to pathway-level dynamics [4]. Consequently, we interpret dependent variables as averages in the conjunctival microenvironment.

The immunopathogenesis of SAC is described by a system of coupled ordinary differential equations (ODEs) denoted by

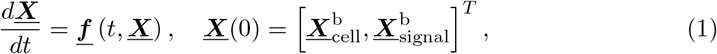

where

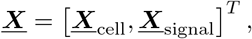

with ***X***_cell_ comprising immune cell populations (in alphabetic order), and ***<u>X</u>***_signal_ comprising the environmental signal (allergens) and immune signals (chemicals and proteins) listed in alphabetic order, and 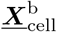 and 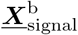 are the immune cell populations and signals baseline values, respectively. The specific components of <u>***X***</u>_cell_

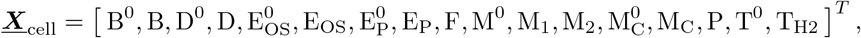

and the components of <u>***X***</u>_signal_ are

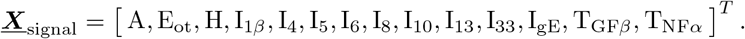

Again, a complete description of each model dependent variable is provided in Table 1. Tables 3 and 4 list baseline values for immune cell populations and immune signals, respectively.

**Table 3.** The immune cell population baseline and homeostatic values.

| Baseline Immune Cell Populations (cells/mL) |  |  | Homeostatic Immune Cell Populations (cells/mL) |  |  |
| --- | --- | --- | --- | --- | --- |
| Variable | Value | Reference | Variable | Fold ($\times X_{ji}^b$) | Reference |
| $B^{0b}$ | (Scale factor) $\times (B^b)$<br>$= (1.0) \times (10^5)$ | [12, 52–54] | $B^{0h}$ | 1.00 | [12, 52–54] |
| $B^b$ | $\frac{\text{Density of B}}{\text{Thickness of conjunctiva}}$<br>$= \frac{5 \text{ cells/mm}^2}{0.05 \text{ mm}} \times \frac{1000 \text{ mm}^3}{\text{mL}}$<br>$\approx 10^5$ | [52–55] | $B^h$ | 1.20 | [51] |
| $D^{0b}$ | (Scale factor) $\times (D^b)$<br>$= (20.0) \times (9.30 \times 10^4)$ | [12, 55, 56] | $D^{0h}$ | 1.00 | [12, 55, 56] |
| $D^b$ | $\frac{\text{Density of D}}{\text{Thickness of conjunctiva}}$<br>$= \frac{4.5 \text{ cells/mm}^2}{0.05 \text{ mm}} \times \frac{1000 \text{ mm}^3}{\text{mL}}$<br>$\approx 9.30 \times 10^4$ | [55, 56] | $D^h$ | 1.00 | [55, 56] |
| $E_{OS}^{0b}$ | (Scale factor) $\times (E_{OS}^b)$<br>$= (1.0) \times (1.60 \times 10^4)$ | [57] | $E_{OS}^{0h}$ | 1.00 | [57] |
| $E_{OS}^b$ | $\frac{\text{Density of } E_{OS}}{\text{Thickness of conjunctiva}}$<br>$= \frac{0.08 \text{ cells/mm}^2}{0.05 \text{ mm}} \times \frac{1000 \text{ mm}^3}{\text{mL}}$<br>$\approx 1.60 \times 10^4$ | [57] | $E_{OS}^h$ | 1.20 | [58, 59] |
| $E_P^{0b}$ | (Scale factor) $\times (E_P^b)$<br>$= (4.8) \times (3.50 \times 10^5)$ | [12, 55, 60] | $E_P^{0h}$ | 1.00 | [12, 55, 60] |
| $E_P^b$ | $\frac{\text{Density of } E_P}{\text{Thickness of conjunctiva}}$<br>$= \frac{17.5 \text{ cells/mm}^2}{0.05 \text{ mm}} \times \frac{1000 \text{ mm}^3}{\text{mL}}$<br>$\approx 3.50 \times 10^5$ | [55, 60] | $E_P^h$ | 1.00 | [55, 60] |
| $F^b$ | $7.20 \times 10^4$ | [61] | $F^h$ | 1.00 | [61] |
| $M^{0b}$ | (Scale factor) $\times (M^b)$<br>$= (20.0) \times (10^5)$ | [12, 54, 62] | $M^{0h}$ | 1.00 | [12, 54, 62] |
| $M_1^b$ | $\frac{\text{Density of } M_1}{\text{Thickness of conjunctiva}}$<br>$= \frac{5 \text{ cells/mm}^2}{0.05 \text{ mm}} \times \frac{1000 \text{ mm}^3}{\text{mL}}$<br>$\approx 10^5$ | [54, 62] | $M_1^h$ | 1.00 | [54, 62] |
| $M_2^b$ | $\frac{\text{Density of } M_2}{\text{Thickness of conjunctiva}}$<br>$= \frac{5 \text{ cells/mm}^2}{0.05 \text{ mm}} \times \frac{1000 \text{ mm}^3}{\text{mL}}$<br>$\approx 10^5$ | [54, 62] | $M_2^h$ | 5.60 | [63] |
| $M_C^{0b}$ | (Scale factor) $\times (M_C^b)$<br>$= (1.0) \times (1.21 \times 10^5)$ | [12, 54] | $M_C^{0h}$ | 1.00 | [12, 54] |
| $M_C^b$ | $\frac{\text{Density of } M_C}{\text{Thickness of conjunctiva}}$<br>$= \frac{6 \text{ cells/mm}^2}{0.05 \text{ mm}} \times \frac{1000 \text{ mm}^3}{\text{mL}}$<br>$\approx 1.21 \times 10^5$ | [54, 55, 57] | $M_C^h$ | 3.00 | [50, 57, 64] |
| $P^b$ | $6.00 \times 10^3$ | [52] | $P^h$ | 7.00 | [51] |
| $T^{0b}$ | (Scale factor) $\times (T_{H2}^b)$<br>$= (4.0) \times (7.85 \times 10^5)$ | [12, 53] | $T^{0h}$ | 1.00 | [12, 53] |
| $T_{H2}^b$ | $\frac{\text{Density of } T_{H2}}{\text{Thickness of conjunctiva}}$<br>$= \frac{39.5 \text{ cells/mm}^2}{0.05 \text{ mm}} \times \frac{1000 \text{ mm}^3}{\text{mL}}$<br>$\approx 7.85 \times 10^5$ | [53] | $T_{H2}^h$ | 1.07 | [53] |

**Table 4.** The immune signal baseline and homeostatic values.

| Baseline Immune Signal Concentrations (pg/mL) |  |  | Homeostatic Immune Signal Concentrations (pg/mL) |  |  |
| --- | --- | --- | --- | --- | --- |
| Variable | Value | Reference | Variable | Fold ($\times X_{ji}^b$) | Reference |
| $E_{ot}^b$ | 6.82 | [65] | $E_{ot}^h$ | 3.00 | [66, 67] |
| $H^b$ | $1.81 \times 10^3$ | [68] | $H^h$ | 3.00 | [69] |
| $I_{1\beta}^b$ | $1.10 \times 10^1$ | [7, 70] | $I_{1\beta}^h$ | 1.00 | [7, 70] |
| $I_4^b$ | 6.00 | [7, 70–72] | $I_4^h$ | 3.00 | [7, 73] |
| $I_5^b$ | 7.00 | [7, 70] | $I_5^h$ | 1.00 | [7, 70] |
| $I_6^b$ | $1.90 \times 10^1$ | [7, 70] | $I_6^h$ | 1.50 | [7, 74] |
| $I_8^b$ | $1.70 \times 10^1$ | [7, 70] | $I_8^h$ | 1.50 | [7, 74, 75] |
| $I_{10}^b$ | 1.00 | [7, 72] | $I_{10}^h$ | 1.00 | [7, 72] |
| $I_{13}^b$ | 2.31 | [7, 72] | $I_{13}^h$ | 1.00 | [7, 72] |
| $I_{33}^b$ | $1.40 \times 10^1$ | [71] | $I_{33}^h$ | 1.00 | [71] |
| $I_{gE}^b$ | $1.00 \times 10^3$ | [76, 77] | $I_{gE}^h$ | 7.50 | [76, 78, 79] |
| $T_{GF\beta}^b$ | $7.17 \times 10^2$ | [65, 80, 81] | $T_{GF\beta}^h$ | 5.50 | [74, 82] |
| $T_{NF\alpha}^b$ | $1.00 \times 10^1$ | [7, 72] | $T_{NF\alpha}^h$ | 1.00 | [7, 72] |

#### 2.2.1 Immune Homeostasis

We assume that in the absence of an environmental stimuli, i.e., allergen antigens, the immune system in the conjunctival microenvironment is in homeostasis, specifically, 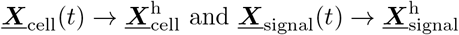 as *t → ∞* when 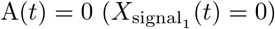. This choice reflects the fact that, in the absence of acute inflammatory stimulation, resident and circulating immune cell pools and signals are maintained by slower homeostatic processes, so the biologically relevant reference state is a nonzero population rather than extinction [47].

We assume the homeostatic values represent inflammatory concentration levels. Consequently, some of the homeostatic values are different from the baseline values. Galli et al. argued that repeated allergen exposure can lead to tissue remodeling and cause the concentrations of the activated cells to keep rising, not returning to their baseline levels even when the allergen are being removed [48]. Fukuda et al. studied tissue remodeling and prolonged inflammation during ocular allergy and showed that the cell numbers in the papillae (conjunctiva) are measured up to 7.3 times higher than their normal subjects [49]. In the experimental study performed by Anderson et al., they demonstrated that the during the immunohistochemical analysis of conjunctival biopsies from patients in and out of pollen season, the lamina propria mast cells remain significantly higher than the healthy population [50]. For IgE-secreting plasmablasts, recent work has shown that these cells can enter a long-lived state in secondary lymphoid tissues after allergen exposure, remaining detectable in the blood for weeks after the triggering stimulus is gone and predicting future disease flares [51]. Tables 3 and 4 list homeostatic values for immune cell populations and signals, respectively.

We assume the naive immune cell and fibroblast cell populations do not change temporal. That is, we assume timescale associated with maintaining homeostasis is much faster then the timescales of acute inflammatory responses. Models of Th2-mediated airway inflammation initiate dynamics from a pre-challenge equilibrium, and models of naive T cell homeostasis justify a constant compartment on acute timescales [83, 84]. Consequently, symbolically, we assume

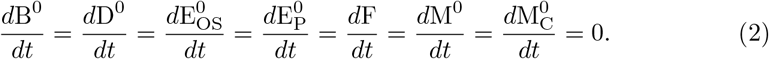

To return the system to homeostasis, a first-order elimination rate of the form

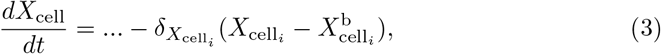

where 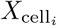 is the *i*-the component of <u>***X***</u>_cell_, 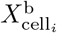 is the baseline value of *i*-the component of <u>***X***</u>_cell_ and 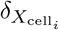 is elimination rate of *i*-the component of <u>***X***</u>_cell_. For the immune cell populations, the elimination rate is taken to be the death rate of the cell.

Differently, for the environmental and immune signals, we assume a first-order elimination rate of the form

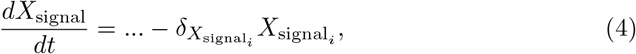

where the elimination rate is zero when the immune signal concentration zero. For the immune signaling vector, the elimination rates are either reported clearance rates (chemical signals) or depletion rates (protein signals) found in the literature. The values of elimination rates are reported in Tables 5 and 6.

**Table 5.** Parameters associated immune cell population dynamics. The first column lists the pathway number from Figure 1 and the last column lists the numbering of the parameter used in Section 4.

| Immune Cell Population Parameters |  |  |  |  |  |  |
| --- | --- | --- | --- | --- | --- | --- |
| Pathway | Parameter | Description | Value | Units | Reference | No. |
| **B cells** |  |  |  |  |  |  |
| 30 | $\lambda_{B,I_4}$ | IL-4 activation rate of B cells | 0.06 | 1/day | [12] | 56 |
| 30 | $K_{I_4,B}$ | Half-saturation IL-4 activation threshold for B cells | 150 | pg/mL | [7, 95] | 40 |
| 4 | $\lambda_{B,A}$ | Allergen activation rate of B cells | 0.06 | 1/day | [12] | 53 |
| 4 | $K_{A,-}$ | Allergen activation half-saturation constant | 100 | pg/mL | [87] | 23 |
| - | $\delta_B$ | Natural death rate of B cells | 0.0141 | 1/day | [96] | 58 |
| **Dendritic cells** |  |  |  |  |  |  |
| 2 | $\lambda_{D,A}$ | Activation rate of dendritic cells by allergen | 0.23 | 1/day | Assumed, Section A.1 | 47 |
| 2 | $K_{A,-}$ | Allergen activation half-saturation constant | 100 | pg/mL | [87] | 23 |
| - | $\delta_D$ | Natural death rate of dendritic cells | 0.231 | 1/day | [96] | 49 |
| **Eosinophil cells** |  |  |  |  |  |  |
| 28 | $\lambda_{E_{OS},I_5}$ | IL-5-mediated eosinophil activation rate | 0.025 | 1/day | [12] | 73 |
| 28 | $K_{I_5,E_{OS}}$ | IL-5 half-saturation constant | 90 | pg/mL | [7] | 64 |
| 13 | $\lambda_{E_{OS},I_{33}}$ | IL-33 activation rate of eosinophils cells | 0.04 | 1/day | [12] | 74 |
| 13 | $K_{I_{33},E_{OS}}$ | Half-saturation IL-33 activation threshold for eosinophils | 970 | pg/mL | [12] | 57 |
| 27 | $\lambda_{E_{OS},E_{ot}}$ | Proliferation rate of eosinophils in the presence of eotaxin | 2.86 | 1/day | Estimated, Section A.2 | 67 |
| 27 | $K_{E_{ot},E_{OS}}$ | Half-saturation activation threshold for eosinophils by eotaxin | 641 | pg/mL | [66] | 61 |
| - | $\delta_{E_{OS}}$ | Natural death rate of eosinophils | 1.67 | 1/day | [12] | 65 |
| **Epithelial cells** |  |  |  |  |  |  |
| 1 | $\lambda_{Ep,A}$ | Activation rate of epithelial cells by allergen | 0.0468 | 1/day | Assumed, Section A.1 | 34 |
| 1 | $K_{A,-}$ | Allergen activation half-saturation constant | 100 | pg/mL | [87] | 23 |
| - | $\delta_{Ep}$ | Natural death rate of epithelial cells | 0.0468 | 1/day | [97] | 43 |
| **M1 macrophages** |  |  |  |  |  |  |
| 3 | $\lambda_{M_1,A}$ | Differentiation rate of naive macrophages into M<sub>1</sub> macrophages | 0.0693 | 1/day | Assumed, Section A.1 | 13 |
| 3 | $K_{A,-}$ | Allergen activation half-saturation constant | 100 | pg/mL | [87] | 23 |
| - | $\delta_{M_1}$ | Natural death rate of M1 macrophages | 0.0693 | 1/day | [98] | 25 |
| **M2 macrophages** |  |  |  |  |  |  |
| 40 | $\lambda_{M_2,I_4}$ | IL-4 proliferation rate by M<sub>2</sub> macrophages | 0.212 | 1/day | Estimated, Section A.2 | 45 |
| 40 | $K_{I_4,M_2}$ | Half-saturation activation of M<sub>2</sub> macrophages by IL-4 | 250 | pg/mL | [7, 99, 100] | 46 |
| - | $\delta_{M_2}$ | Natural death rate of M2 macrophages | 0.0693 | 1/day | [98] | 55 |
| **Mast cells** |  |  |  |  |  |  |
| 32 | $\lambda_{M_C,I_{gE}}$ | Mast-cells degranulation in the presence IgE antibody | 5.0941 | 1/day | Estimated, Section A.2 | 17 |
| 32 | $K_{I_{gE},M_C}$ | Half-saturation of IgE | 20000 | pg/mL | [12] | 20 |
| 31 | $k_{I_{10},M_C}$ | IL-10 concentration inhibiting mast cell activation | 100 | pg/mL | Estimated, Section A.3 | 59 |
| - | $\delta_{M_C}$ | Natural death rate of mast cells | 0.693 | 1/day | [101] | 19 |
| Pathway | Parameter | Description | Value | Units | Reference | No. |
| Plasma cells |  |  |  |  |  |  |
| 5 | $\lambda_{P,A}$ | Plasma cell differentiation rate from B cells | 0.06 | 1/day | [12] | 30 |
| 5 | $K_{A,-}$ | Allergen activation half-saturation constant | 100 | pg/mL | [87] | 23 |
| 37 | $\lambda_{P,I_4}$ | Plasma cell activation rate by IL-4 | 0.06 | 1/day | [12] | 41 |
| 37 | $K_{I_4,B}$ | Half-saturation IL-4 activation threshold for B cells | 150 | pg/mL | [7, 95] | 40 |
| - | $\delta_P$ | Natural death rate of plasma cells | 0.02 | 1/day | [102] | 48 |
| Th2 cells |  |  |  |  |  |  |
| 12 | $\lambda_{T_{H2},I_{1\beta}}$ | Th2 proliferation rate induced by IL-1β | 0.01 | 1/day | Estimated, Section A.2 | 50 |
| 12 | $K_{I_{1\beta},T_{H2}}$ | IL-1β half-saturation constant | 120 | pg/mL | [7, 103] | 51 |
| 10 | $\lambda_{T_{H2},I_6}$ | Th2 proliferation rate induced by IL-6 | 0.159 | 1/day | Estimated, Section A.2 | 36 |
| 10 | $K_{I_6,T_{H2}}$ | IL-6 half-saturation constant | 353 | pg/mL | [7, 104] | 37 |
| 11 | $\lambda_{T_{H2},I_{33}}$ | Th2 proliferation rate induced by IL-33 | 0.0605 | 1/day | Estimated, Section A.2 | 76 |
| 14 | $\lambda_{T_{H2},I_4}$ | Th2 proliferation rate induced by IL-4 | 0.0447 | 1/day | Estimated, Section A.2 | 39 |
| 14 | $K_{I_4,T_{H2}}$ | IL-4 half-saturation constant | 250 | pg/mL | [7, 99, 100] | 42 |
| 11 | $K_{I_{33},T_{H2}}$ | Half-saturation IL-33 activation threshold for Th2 cells | 1000 | pg/mL | [12] | 77 |
| 15 | $k_{T_{GF\beta},T_{H2}}$ | TGF-β inhibition threshold for Th2 activity | 444 | pg/mL | Estimated, Section A.4 | 28 |
| - | $\delta_{T_{H2}}$ | Natural death rate of Th2 cells | 0.1 | 1/day | [96] | 32 |

**Table 6.** Parameters associated immune cell population dynamics. The first column lists the pathway number from Figure 1 and the last column lists the numbering of the parameter used in Section 4.

| Immune Signal Concentration Parameters |  |  |  |  |  |  |
| --- | --- | --- | --- | --- | --- | --- |
| Pathway | Parameter | Description | Value | Units | Reference | No. |
| **Eotaxins** |  |  |  |  |  |  |
| 24 | $\psi_{E_{ot}, F}$ | Production rate of eotaxin by fibroblasts | $5.17 \times 10^{-4}$ | pg/cell/day | Estimated, Section A.5 | 66 |
| 24 | $K_{I_4, E_{ot}}$ | IL-4 half-saturation constant | $2.50 \times 10^2$ | pg/mL | [7, 99, 100] | 60 |
| - | $\delta_{E_{ot}}$ | Depletion rate of eotaxin | $1.00 \times 10^{-1}$ | 1/day | [105] | 68 |
| **Histamines** |  |  |  |  |  |  |
| 36 | $\psi_{H, M_C}$ | Histamine production rate by mast cells | $1.234 \times 10^{-1}$ | pg/cell/day | Estimated, Section A.5 | 72 |
| - | $\delta_H$ | Clearance rate of histamine | 8.32 | 1/day | [106] | 69 |
| **IL-1β** |  |  |  |  |  |  |
| 29 | $\psi_{I_{1\beta}, M_1}$ | IL-1β production rate by M<sub>1</sub> macrophages | $7.35 \times 10^{-4}$ | pg/cell/day | Estimated, Section A.5 | 4 |
| - | $\delta_{I_{1\beta}}$ | Depletion rate of IL-1β | 6.65 | 1/day | [107] | 9 |
| **IL-4** |  |  |  |  |  |  |
| 22 | $\psi_{I_4, E_{OS}}$ | IL-4 production rate by eosinophils | $5.00 \times 10^{-6}$ | pg/cell/day | [12] | 63 |
| 23 | $\psi_{I_4, M_C}$ | IL-4 production rate by mast cells | $3.52 \times 10^{-4}$ | pg/cell/day | Estimated, Section A.5 | 2 |
| 20 | $\psi_{I_4, T_{H2}}$ | IL-4 production rate by Th2 cells | $3.25 \times 10^{-5}$ | pg/cell/day | [12] | 52 |
| 20 | $K_{I_{13}, I_4}$ | Half-saturation constant of IL-13 | $3.50 \times 10^1$ | pg/mL | [7] | 54 |
| 21 | $k_{I_{10}, I_4}$ | IL-10 inhibition threshold for IL-4 production | $1.00 \times 10^2$ | pg/mL | Estimated, Section A.3 | 62 |
| - | $\delta_{I_4}$ | Depletion rate of IL-4 | 6.9315 | 1/day | [12] | 1 |
| **IL-5** |  |  |  |  |  |  |
| 18 | $\psi_{I_5, T_{H2}}$ | IL-5 production rate by Th2 cells | $5.90 \times 10^{-5}$ | pg/cell/day | [12] | 5 |
| - | $\delta_{I_5}$ | Depletion rate of IL-5 | 6.9315 | 1/day | [12] | 6 |
| **IL-6** |  |  |  |  |  |  |
| 6 | $\psi_{I_6, D}$ | IL-6 production rate by dendritic cells | $1.39 \times 10^{-4}$ | pg/cell/day | Estimated, Section A.5 | 27 |
| 7 | $\psi_{I_6, M_1}$ | IL-6 production rate by M1 macrophages | $1.39 \times 10^{-4}$ | pg/cell/day | Estimated, Section A.5 | 16 |
| - | $\delta_{I_6}$ | Depletion rate of IL-6 | 1.07 | 1/day | [108] | 11 |
| **IL-8** |  |  |  |  |  |  |
| 25 | $\psi_{I_8, F}$ | IL-8 production rate by fibroblasts | $1.7 \times 10^{-3}$ | pg/cell/day | Estimated, Section A.5 | 33 |
| 25 | $K_{I_4, I_8}$ | Half-saturation IL-4 threshold constant for proliferation of IL-8 | $1.00 \times 10^2$ | pg/mL | [109, 110] | 35 |
| 26 | $\psi_{I_8, F}$ | IL-8 production rate by fibroblasts | $1.7 \times 10^{-3}$ | pg/cell/day | Estimated, Section A.5 | 33 |
| 26 | $K_{T_{NF\alpha}, I_8}$ | Half-saturation constant of TNF-α for proliferation of IL-8 | $1.08 \times 10^1$ | pg/mL | [7, 111] | 44 |
| 9 | $\psi_{I_8, E_P}$ | IL-8 production rate by epithelial cells | $5.29 \times 10^{-5}$ | pg/cell/day | Estimated, Section A.5 | 15 |
| - | $\delta_{I_8}$ | Depletion rate of IL-8 | 1.109 | 1/day | [112] | 12 |
| Pathway | Parameter | Description | Value | Units | Reference | No. |
| **IL-10** |  |  |  |  |  |  |
| 16 | $\psi_{I_{10}, T_{H2}}$ | IL-10 production rate by Th2 cells | $4.97 \times 10^{-6}$ | pg/cell/day | Estimated, Section A.5 | 14 |
| 17 | $\psi_{I_{10}, M_2}$ | IL-10 production rate by M2 macrophages | $4.97 \times 10^{-6}$ | pg/cell/day | Estimated, Section A.5 | 38 |
| - | $\delta_{I_{10}}$ | Depletion rate of IL-10 | 6.9315 | 1/day | [12] | 7 |
| **IL-13** |  |  |  |  |  |  |
| 19 | $\psi_{I_{13}, T_{H2}}$ | IL-13 production rate by Th2 cells | $1.91 \times 10^{-5}$ | pg/cell/day | Estimated, Section A.5 | 10 |
| - | $\delta_{I_{13}}$ | Depletion rate of IL-13 | 6.9315 | 1/day | [12] | 8 |
| **IL-33** |  |  |  |  |  |  |
| 8 | $\psi_{I_{33}, E_P}$ | IL-33 production rate by epithelial cells | $2.77 \times 10^{-4}$ | pg/cell/day | [12] | 71 |
| - | $\delta_{I_{33}}$ | Depletion rate of IL-33 | 6.93 | 1/day | [12] | 70 |
| **IgE** |  |  |  |  |  |  |
| 38 | $\psi_{I_{IgE}, P}$ | IgE production rate by plasma cells | $5.84 \times 10^{-2}$ | pg/cell/day | Estimated, Section A.5 | 21 |
| - | $\delta_{I_{IgE}}$ | Clearance rate of IgE | $3.46 \times 10^{-1}$ | 1/day | [101] | 24 |
| **TGF-β** |  |  |  |  |  |  |
| 39 | $\psi_{T_{GF\beta}, M_2}$ | TGF-β production rate by M2 macrophages | 3.493 | pg/cell/day | Estimated, Section A.5 | 29 |
| - | $\delta_{T_{GF\beta}}$ | Depletion rate of TGF-β | 499 | 1/day | [113] | 31 |
| **TNF-α** |  |  |  |  |  |  |
| 33 | $\psi_{T_{NF\alpha}, M_1}$ | TNF-α production rate by M1 macrophages | $1.06 \times 10^{-2}$ | pg/cell/day | Estimated, Section A.5 | 18 |
| 35 | $k_{I_{10}, T_{NF\alpha}}$ | IL-10 inhibition threshold for TNF-α production | $1.00 \times 10^2$ | pg/mL | Estimated, Section A.3 | 75 |
| 34 | $\psi_{T_{NF\alpha}, M_C}$ | TNF-α production rate by mast cells | $1.06 \times 10^{-3}$ | pg/cell/day | Estimated, Section A.5 | 22 |
| - | $\delta_{T_{NF\alpha}}$ | Depletion rate of TNF-α | 47.53 | 1/day | [114] | 3 |

#### 2.2.2 Immune Response

Allergen antigens active an immune response. The immunopathological mechanisms in SAC can be characterized broadly as either a (i) **change-of-state** or (ii) **production**. In general, immune cells can both change their state and produce immune signals, and the immune signals can activate or inhibit these mechanisms. Next, we describe how we quantify each mechanism.

##### Activated immune cell change-of-state without inhibition

Each immune cell change-of-state is activated by a signal. We model the activation of the change-of-state of 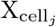 by the *i*-th signal 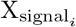 using a Hill function of the form

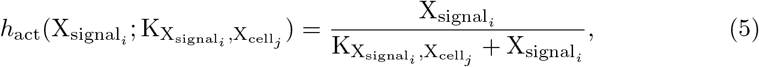

where 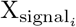 is concentration the activation signal and 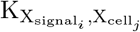 is the half-saturation concentration of 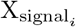 for activating 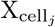. Note that if half-saturation concentration of 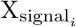 is the same for all cells, then we denote by 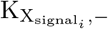. This formulation captures essential biological features of immune activation, including threshold behavior, and saturation [85]. In immune and infection modeling, sigmoidal activation terms are commonly used to describe cytokine-mediated immune cell activation and pathogen-driven inflammatory responses, where increasing stimulus leads to diminishing marginal effects at high concentrations [86].

The nonlinear activation function characterizes the change-of-state of immune cells; more specifically, antigen-presenting cell activation by allergens (pathways 1-4), cell differentiation activated by allergens (pathway 5), cytokine-dependent differentiation of cells (pathway 37), and cell activation after inflammatory stimulation (pathways 13, 27, 28, 30, and 40). Mathematically, the change-of-state of immune cell X_cell_*i* to X_cell_*j* activated by X_signal_*k* is modeled by

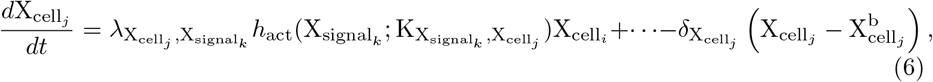

where 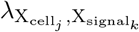 is change-of-state rate, 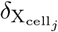 is the first-order elimination rate, and 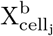 is the baseline values reported in Table 3. In general, we assume the rate of change of the *j*-th immune cell is proportional to the concentration of the *i*-th immune cell, where the proportional constant is modified by nonlinear activation function due to signal 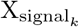.

##### Activated immune cell change-of-state pathways with inhibition

An immune cell change-of-state can be both activated by a signal and also inhibited by a signal. More specifically, we assume the regulatory cytokines IL-10 and TGF-*β* are inhibitory modulators of inflammatory processes. We model the inhibition using the Hill-type suppression function of the form

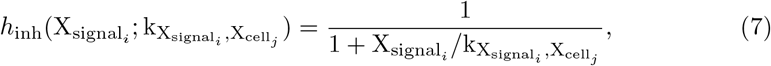

where 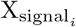 represents the concentration of the inhibitory cytokine and 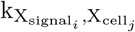 is the half inhibition concentration. This functional form reflects the biological role of regulatory cytokines as modulators rather than as absolute suppressors of immune activity. Experimental and modeling studies of immune regulation indicate that inhibitory signaling attenuates pro-inflammatory responses in a graded, concentration-dependent manner rather than enforcing complete shutdown. Consequently, inhibitory Hill functions are commonly used in mechanistic immune and gene-regulatory models to represent negative feedback while preserving residual immune responsiveness [85, 86].

In the governing equations, the nonlinear inhibition function multiplies immune cell change-of-state rates allowing regulatory effects to scale with the magnitude of the underlying inflammatory signal. Mathematically, the change-of-state of immune cell 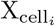 to 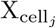 activated by 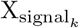 and inhibited by 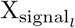 is modeled by

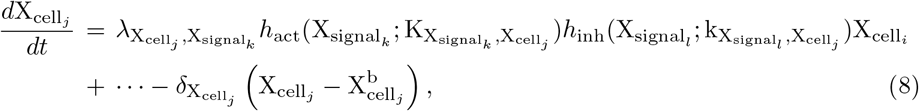

where 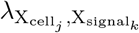is change-of-state rate, 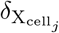 is the elimination rate, and 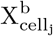 is the baseline value reported in Table 3.

##### Immune signal production pathways without and with inhibition

Mature immune cells produce immune signals. We assume the immune signal production is proportional mature immune cell population. Without inhibition, the production of 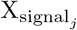 by 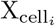 is given by

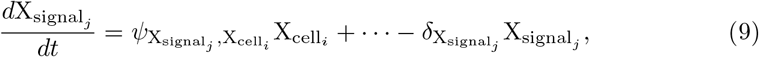

where 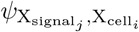 is the production rate, 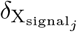 is the elimination rate. If the production is inhibited by 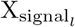 then the the production of 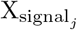 by 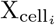 is given by

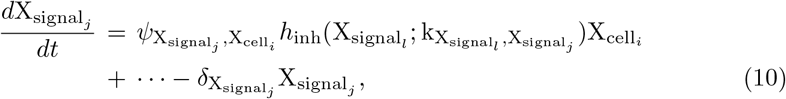

where 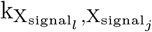 is the concentration of 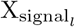 that inhibits the production rate by 50%. This structure captures the biologically observed balance between immune activation and regulation.

##### Activated immune signal production pathways without and with inhibition

Immune cells, specifically, Th2 cells and fibroblasts, produce immune signals when activated by a different immune signal. The production of immune signal 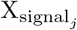 by 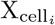 activated by 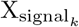 without inhibition is given by

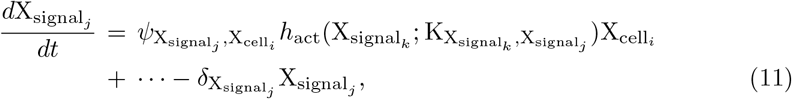

where 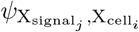 is the production rate, 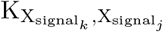 is the half-saturation concentration of 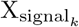 for activating 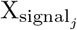, and 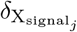is the elimination rate.

If 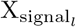 is inhibiting the production, then the production of immune signal 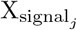 by 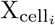 activated by 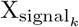 without inhibition is given by

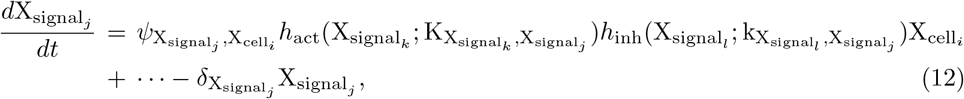

where 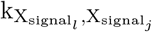 is the half-inhibition concentration.

##### Allergen antigen dynamics

The allergen antigen dynamics follows the same pattern as ragweed pollen concentration data presented in Bonini et al. [87]. Figure 2 (blue, left axis) plots the digitized data from Bonini et al. of daily ragweed pollen grain concentrations at different locations in Italy over a 63-day pollen season.

**Fig 2.**
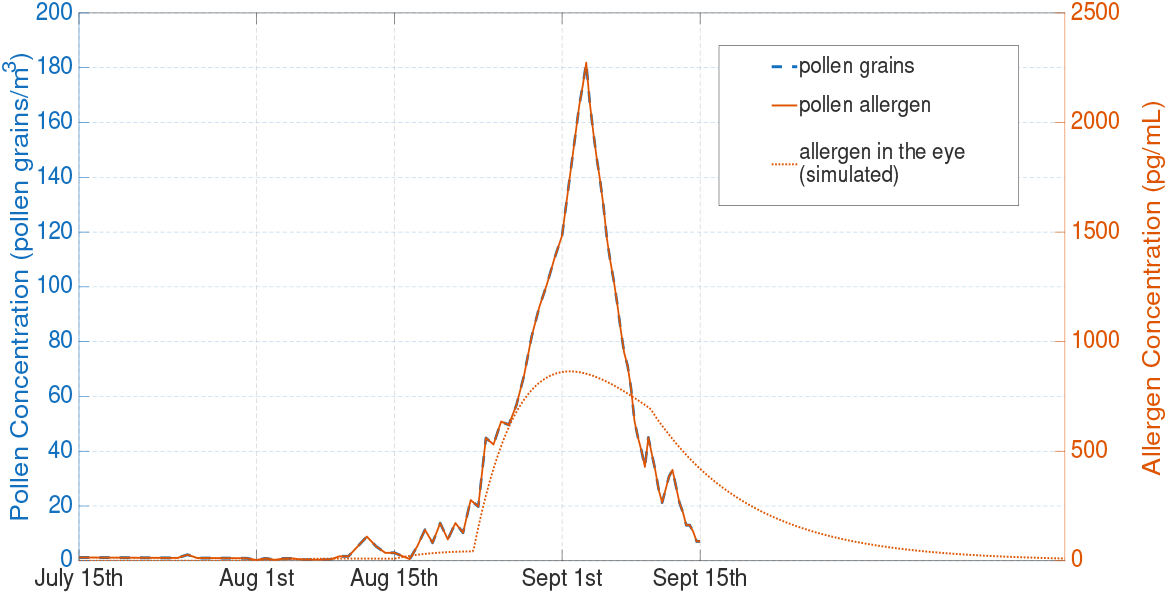
Seasonal pollen exposure and modeled ocular allergen antigen in the tear. Measured ragweed pollen grain concentrations (blue, left axis). Data is used to generate a continuous allergen input function representing allergen concentration within the ocular surface compartment (orange, right axis), which is triggers the immune response in the model.

The major allergen in ragweed pollen is referred to as Amb a 1 and is responsible for over 90% of allergic responses in sensitive individuals [88]. In some studies, ragweed allergen has been recorded to range from 4.57 pg per pollen grain for aged pollen to 16.5 pg per pollen grain [89]. In our work, we assume the airborne pollen grain concentration is proportional to the allergen antigens present in the tears. Therefore, we are using the conversion factor of 1 pollen grain/m^3^ = 12.5 pg/mL Amb a 1 representing a typical intermediate value for ambient air conditions [89]. This conversion factor is a biologically plausible value for estimating allergen exposure from Bonini et al.’s study [87]. The converted values are graphed as orange line (right axis) in Figure 2.

We model the allergen antigen concentration dynamics in the conjunctivitis microenvironment as follows:

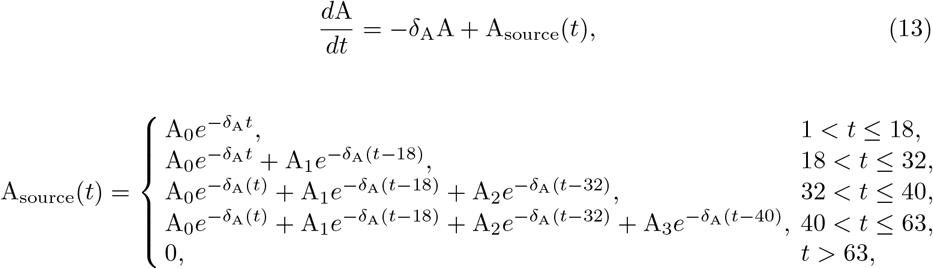

where *δ*_A_ = 0.1 1/day is the clearance rate of allergen, and A_0_ = 0.11 pg/mL, A_1_ = 2.70 pg/mL, A_2_ = 10.1 pg/mL, and A_3_ = 225.8 pg/mL found using the Bonini et al.’s experimental data but scaled by 10% [87]. We scale by 10% to represent transfer of airborne allergen to the ocular surface and its dissolution into the tear film. This represent a biologically plausible allergen concentration within the ocular surface compartment. Popov et al. demonstrates that the ocular mucus layer acts as a significant barrier, with only a small fraction of particulate systems reaching anterior ocular tissues [90]. This is consistent with ophthalmic drug delivery literature reporting that approximately 5 − 10% fraction of administered topical dose reaches ocular target tissues due to rapid tear turnover and mucosal clearance mechanisms [90, 91].

The source term for allergen antigens is modeled as a sum of exponentially decaying functions to represent repeated, time-localized pollen exposure events during the ragweed season. This formulation is motivated by pharmacokinetic modeling principles, in which exponential decay is commonly used to describe first-order clearance processes following bolus-like inputs [92, 93]. Each exponential term represents a discrete pollen deposition event followed by first-order depletion of antigen due to tear turnover, blinking, and epithelial clearance. The piecewise structure allows cumulative exposure during periods of repeated environmental contact [16, 94]. After the pollen season ends, the source term is set to zero, reflecting the absence of new allergen input and complete clearance of residual antigens.

The measured pollen allergen is used to construct a continuous allergen input function. This airborne allergen input is scaled by 10% to represent transfer of airborne allergen to the ocular surface and its dissolution into the tear film. This represent a biologically plausible allergen concentration within the ocular surface compartment. Popov et al. demonstrates that the ocular mucus layer acts as a significant barrier, with only a small fraction of particulate systems reaching anterior ocular tissues [90]. This is consistent with ophthalmic drug delivery literature reporting that approximately 5 *−* 10% fraction of administered topical dose reaches ocular target tissues due to rapid tear turnover and mucosal clearance mechanisms [90, 91]. We assume a similar phenomena take place with allergen penetration on the ocular surface. Consequently, the allergen concentration in the eye is the orange dashed line in Figure 2.

#### 2.2.3 Model Equations

Our complete mathematical model is written here. First, the system of the ordinary differential equations for <u>***X***</u>_cell_:

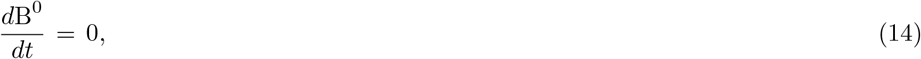

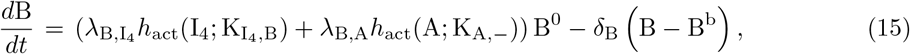

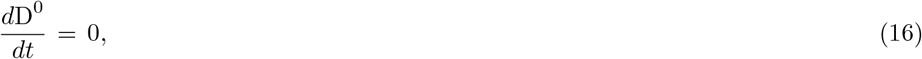

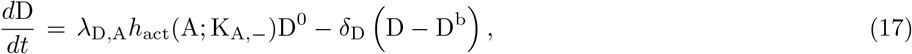

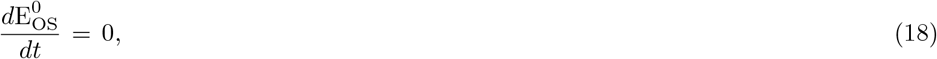

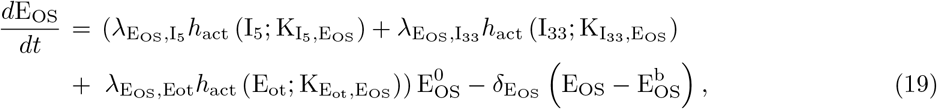

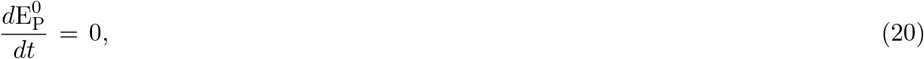

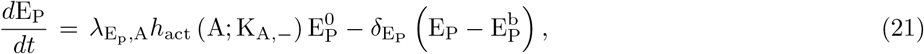

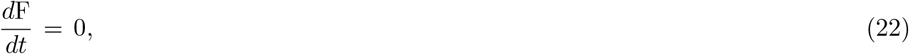

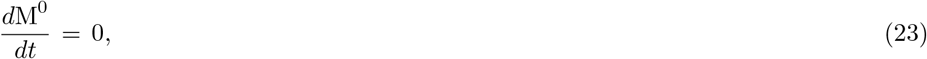

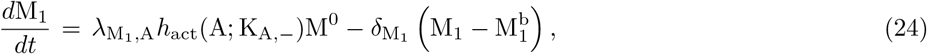

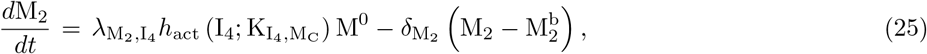

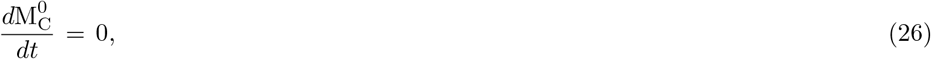

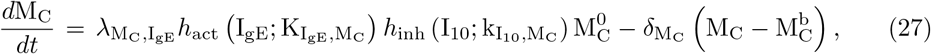

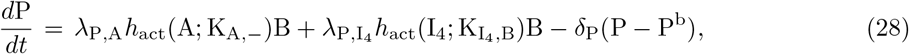

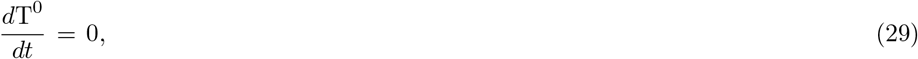

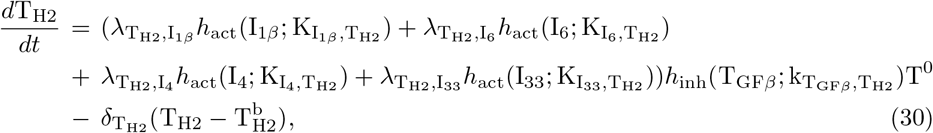

where 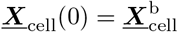. A complete parameter list, in the order they appeared above, is given in Table 5. Parameter values were found either in the literature or estimated from the imposed homeostatic values. Details on how parameter values listed as “Assumed” or “Estimated” can be found in Section A.

The system of differential equations for ***X***_signal_:

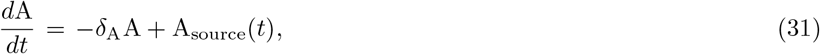

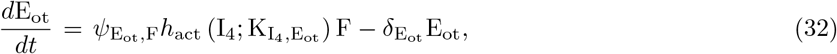

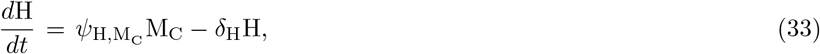

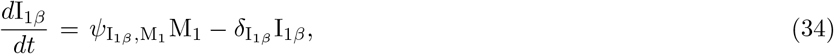

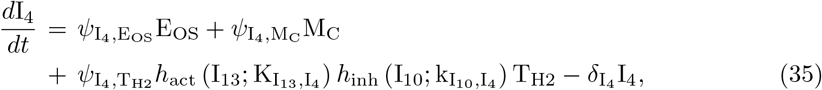

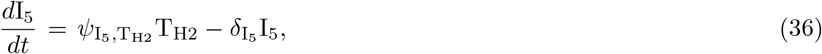

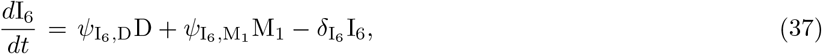

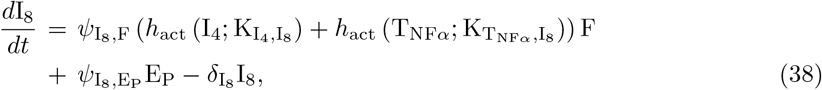

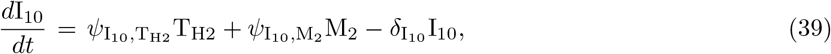

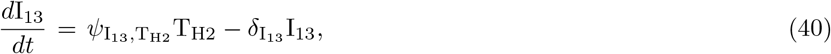

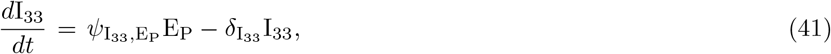

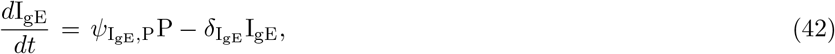

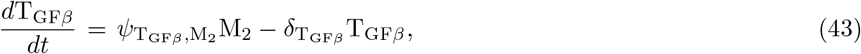

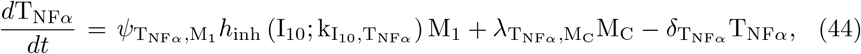

where 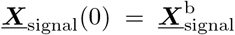. A complete parameter list, in the order they appear above, is given in Table 6. Parameter values were found either in the literature or estimated from the imposed homeostatic values. Details on how parameter values listed as “Assumed” or “Estimated” can be found in Section A.

## 3. Methods

### 3.1 Model Simulations

We approximated the system of ordinary differential equations, stated generally in Equation (1), using MATLAB ode15s. The numerical algorithm was verified by confirming the system evolved to the imposed homeostatic values.

### 3.2 Model Calibration

We calibrated our model to experimental data. Here, we first introduce the experimental data and then we provide an overview of our multi-step model calibration approach.

#### Experimental data

Leonardi et al. measured tear cytokine concentrations in 12 patients with SAC. Tear samples were collected during the pollen season without specification of exact sampling dates [7]. The collected tears were assayed using multiplex bead analysis for cytokines and chemokines including IL-1*β*, IL-4, IL-5, IL-6, IL-8, IL-10, IL-13, and TNF-*α*. Table 7 reports the summary statistics of the experimental data used for model calibration.

**Table 7.** Summary statistics of the experimental measured immune signal concentrations taken from [7] and used in model calibration.

| Immune Signal<br>$k$ | Median (pg/mL)<br>$M^k$ | 1st Quantile (pg/mL)<br>$Q_1^k$ | 3rd Quantile (pg/mL)<br>$Q_3^k$ |
| --- | --- | --- | --- |
| IL-1β | 86.05 | 45.0 | 165.9 |
| IL-4 | 151.6 | 17.1 | 306.0 |
| IL-5 | 80.6 | 38.5 | 143.4 |
| IL-6 | 138.0 | 117.8 | 225.9 |
| IL-8 | 212.5 | 151.6 | 338.6 |
| IL-10 | 4.15 | 2 | 8.6 |
| IL-13 | 13.3 | 10.7 | 32 |
| TNF-α | 13.9 | 10 | 17 |

Bonini et al. not only measured pollen levels in Italy, but they also tracked patient conjunctivitis clinical diary scores [87]. Symptom scores were recorded between July 16th and September 15th (compare timeline with the measured pollen season in Figure 2). The conjunctivitis clinical diary symptom score provides a daily measure of patient-reported symptoms. Individual symptoms are rated on a four-point scale, where 0 denotes no symptoms and 3 denotes severe symptoms. Reported symptoms include ocular itching, throat itching, and burning of the eyes, among others, and are aggregated into a total symptom score ranging from 0 to 24. This composite score comprises contributions from nasal (0–9), ocular (0–6), and bronchial (0–9) symptom subscores [87]. We compare our model to the total composite score observed in a patient group (42 patients) for allergic conjunctivitis [87].

#### Overview of model calibration

A schematic of the multi-step model calibration process used is shown in Figure 3. In the first two steps, we selected “influential” parameters to calibrated using uncertainty quantification [115, 116]. Due to the computational costs of our global sensitivity analysis technique, we first conducted local sensitivity analysis to identify parameters with minimal influence on experimentally-measured model outputs. Parameters that exhibited low sensitivity were subsequently filtered from further analysis. Global sensitivity analysis was then performed to identify the “most influential” parameters to be fit to the experimental data. In the third step, the structural identifiability of the selected parameters was assessed to determine whether unique parameter values could be inferred from the available model outputs [115]. Finally, parameter estimation was carried out by combining regression-based fitting with numerical optimization to calibrate model parameters against experimental data. Next, in Sections 3.2.1-3.2.3, we explain the details of each method implemented.

**Fig 3.**
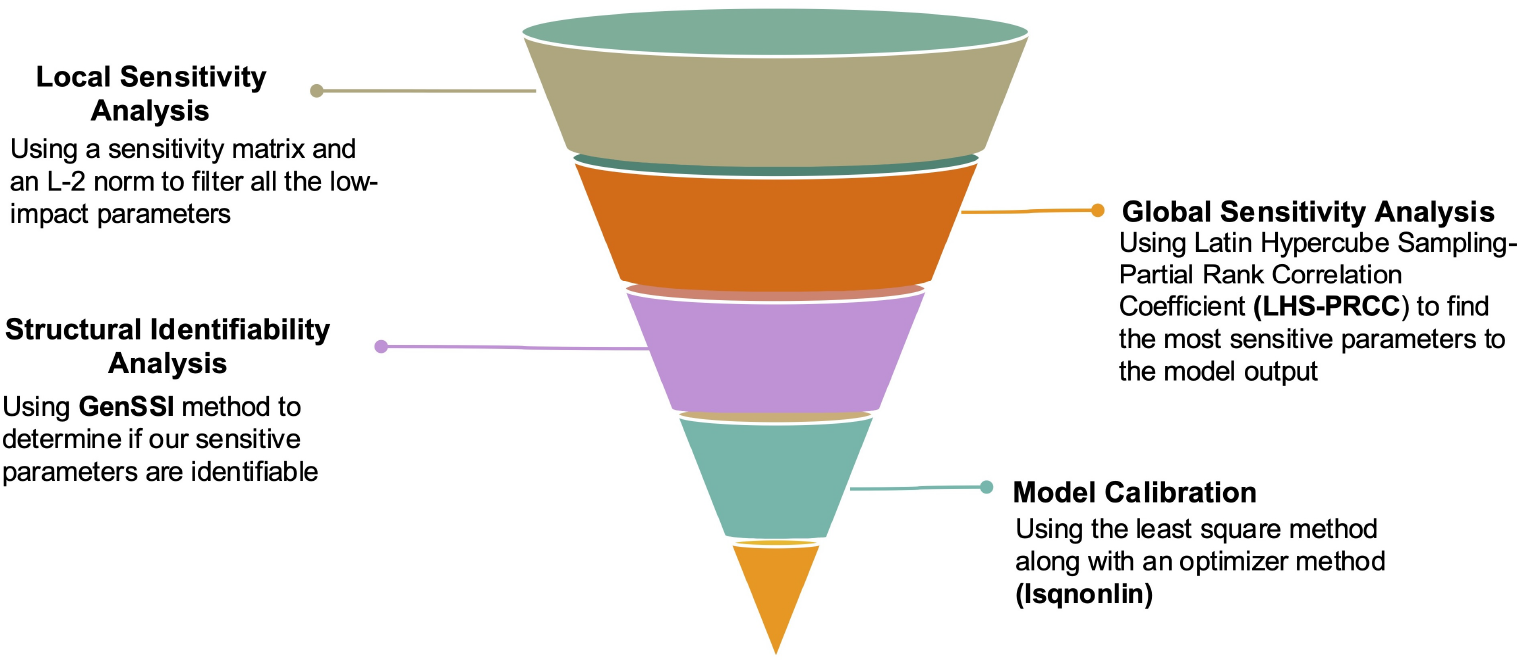
Schematic representing the steps to calibrate the model.

#### 3.2.1 Parameter Selection

We selected parameters for model calibration using a two-pronged sensitivity analysis approach.

##### Local Sensitivity Analysis

We first constructed a sensitivity matrix for the experimentally measurable outputs. Let 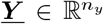 denoted the measurable immune signal outputs, where

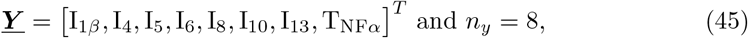

a subset of <u>***X***</u>_signal_. Similar to prior works, we built a relative sensitivity matrix 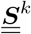 for the *k*-th measured output as follows:

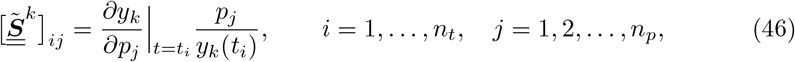

where *pj* is the *j*-th indexed parameter, 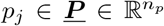 (a complete parameter list written in Table 8) and time *ti* corresponds to the days of pollen season (*ti* = *i* day, where *i* = 32 corresponds to August 15th and *i* = 63 to September 15th, see Figure 2) [116, 117]. This step ensured that we can compare the influence of parameters of different units and magnitudes on the different model outputs *yk*.

**Table 8:** Local sensitivity indices 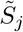 for model parameters. Parameters are ranked in descending order of influence.

| No. | Parameter | $\tilde{S}_j$ | No. | Parameter | $\tilde{S}_j$ |
| --- | --- | --- | --- | --- | --- |
| 1 | $\delta_{I_4}$ | 0.4085 | 41 | $\lambda_{P,I_4}$ | 0.0422 |
| 2 | $\psi_{I_4,M_C}$ | 0.4057 | 42 | $K_{I_4,T_{H2}}$ | 0.0421 |
| 3 | $\delta_{T_{NF\alpha}}$ | 0.3519 | 43 | $\delta_{E_P}$ | 0.0356 |
| 4 | $\psi_{I_{1\beta},M_1}$ | 0.3511 | 44 | $K_{T_{NF\alpha},I_8}$ | 0.0320 |
| 5 | $\psi_{I_5,T_{H2}}$ | 0.3507 | 45 | $\lambda_{M_2,I_4}$ | 0.0306 |
| 6 | $\delta_{I_5}$ | 0.3504 | 46 | $K_{I_4,M_2}$ | 0.0298 |
| 7 | $\delta_{I_{10}}$ | 0.3502 | 47 | $\lambda_{D,A}$ | 0.0271 |
| 8 | $\delta_{I_{13}}$ | 0.3500 | 48 | $\delta_P$ | 0.0211 |
| 9 | $\delta_{I_{1\beta}}$ | 0.3490 | 49 | $\delta_D$ | 0.0203 |
| 10 | $\psi_{I_{13},T_{H2}}$ | 0.3488 | 50 | $\lambda_{T_{H2},I_{1\beta}}$ | 0.0202 |
| 11 | $\delta_{I_6}$ | 0.3481 | 51 | $K_{I_{1\beta},T_{H2}}$ | 0.0147 |
| 12 | $\delta_{I_8}$ | 0.3428 | 52 | $\psi_{I_4,T_{H2}}$ | 0.0109 |
| 13 | $\lambda_{M_1,A}$ | 0.3207 | 53 | $\lambda_{B,A}$ | 0.0104 |
| 14 | $\psi_{I_{10},T_{H2}}$ | 0.3073 | 54 | $K_{I_{13},I_4}$ | 0.0102 |
| 15 | $\psi_{I_8,E_P}$ | 0.2715 | 55 | $\delta_{M_2}$ | 0.0091 |
| 16 | $\psi_{I_6,M_1}$ | 0.2465 | 56 | $\lambda_{B,I_4}$ | 0.0039 |
| 17 | $\lambda_{M_C,I_{gE}}$ | 0.2188 | 57 | $K_{I_{33},E_{OS}}$ | 0.0035 |
| 18 | $\psi_{T_{NF\alpha},M_1}$ | 0.2085 | 58 | $\delta_B$ | 0.0019 |
| 19 | $\delta_{M_C}$ | 0.2057 | 59 | $k_{I_{10},M_C}$ | 0.0018 |
| 20 | $K_{I_{gE},M_C}$ | 0.1791 | 60 | $K_{I_4,E_{ot}}$ | $6.48 \times 10^{-4}$ |
| 21 | $\psi_{I_{gE},P}$ | 0.1732 | 61 | $K_{E_{ot},E_{OS}}$ | $5.69 \times 10^{-4}$ |
| 22 | $\psi_{T_{NF\alpha},M_C}$ | 0.1599 | 62 | $k_{I_{10},I_4}$ | $4.91 \times 10^{-4}$ |
| 23 | $K_{A,-}$ | 0.1596 | 63 | $\psi_{I_4, E_{OS}}$ | $4.88 \times 10^{-4}$ |
| 24 | $\delta_{I_{gE}}$ | 0.1523 | 64 | $K_{I_5, E_{OS}}$ | $1.62 \times 10^{-4}$ |
| 25 | $\delta_{M_1}$ | 0.1429 | 65 | $\delta_{E_{OS}}$ | $6.57 \times 10^{-5}$ |
| 26 | $\delta_A$ | 0.1292 | 66 | $\psi_{E_{ot}, F}$ | $6.50 \times 10^{-5}$ |
| 27 | $\psi_{I_6, D}$ | 0.1271 | 67 | $\lambda_{E_{OS}, E_{ot}}$ | $5.29 \times 10^{-5}$ |
| 28 | $k_{T_{GF\beta}, T_{H2}}$ | 0.08114 | 68 | $\delta_{E_{ot}}$ | $4.54 \times 10^{-5}$ |
| 29 | $\psi_{T_{GF\beta}, M_2}$ | 0.08110 | 69 | $\delta_H$ | $3.26 \times 10^{-5}$ |
| 30 | $\lambda_{P, A}$ | 0.0809 | 70 | $\delta_{I_{33}}$ | $2.05 \times 10^{-5}$ |
| 31 | $\delta_{T_{GF\beta}}$ | 0.0807 | 71 | $\psi_{I_{33}, E_P}$ | $8.57 \times 10^{-6}$ |
| 32 | $\delta_{T_{H2}}$ | 0.0799 | 72 | $\psi_{H, M_C}$ | $5.56 \times 10^{-7}$ |
| 33 | $\lambda_{I_8, F}$ | 0.0798 | 73 | $\lambda_{E_{OS}, I_5}$ | $5.45 \times 10^{-7}$ |
| 34 | $\lambda_{E_P, A}$ | 0.0745 | 74 | $\lambda_{E_{OS}, I_{33}}$ | $2.36 \times 10^{-7}$ |
| 35 | $K_{I_4, I_8}$ | 0.0697 | 75 | $k_{I_{10}, T_{NF\alpha}}$ | $\approx 0$ |
| 36 | $\lambda_{T_{H2}, I_6}$ | 0.0676 | 76 | $\lambda_{T_{H2}, I_{33}}$ | $\approx 0$ |
| 37 | $K_{I_6, T_{H2}}$ | 0.0523 | 77 | $K_{I_{33}, T_{H2}}$ | $\approx 0$ |
| 38 | $\psi_{I_{10}, M_2}$ | 0.0469 | | | |
| 39 | $\lambda_{T_{H2}, I_4}$ | 0.0435 | | | |
| 40 | $K_{I_4, B}$ | 0.0424 | | | |

The partial derivative was approximated using a forward finite difference scheme:

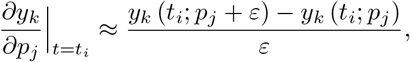

where 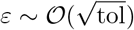 and tol is defined as the relative tolerance of the solver used. In our case, we used ode15s in MATLAB where we set the relative and absolute tolerances such that tol = 10^*−*16^. The choice of tol was due to our smallest parameter having an order of 10^*−*7^.

Using the *k*-th relative sensitivity matrix, we constructed the relative sensitivity matrix 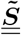 given by

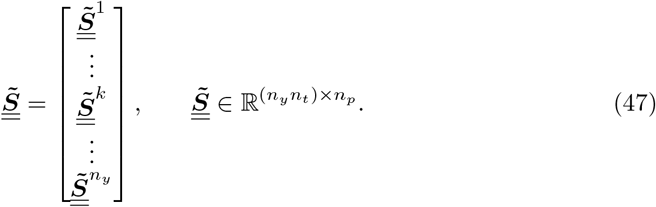

A scalar sensitivity index 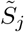 of all measurable outputs over the entire pollen season to each parameter *pj* was computed by taking the *L*2-norm of the *j*-the column of 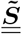 and then normalized by the total number of observations:

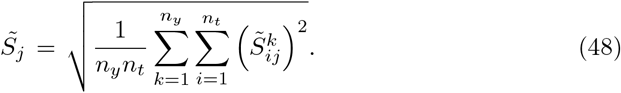

We identified parameter *pj* as locally influential, if

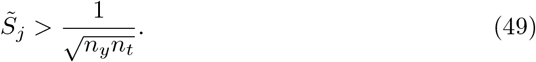

This threshold corresponds to the sensitivity index of a hypothetical parameter whose relative sensitivity value equals exactly 1 (relative output change is equal to the relative parameter change) for one output and at a single time point. Consequently,we identified a subset of parameters 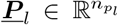, where *np < np*, that are locally influential.

##### Global Sensitivity Analysis

Next, we assessed the global influence of the identified locally influential parameters. A global sensitivity analysis was performed using a Latin hypercube sampling–based partial rank correlation coefficient (LHS–PRCC) approach [118]. Unlike local sensitivity analysis, which evaluated parameter effects around a single, nominal parameter set, the LHS–PRCC framework quantified monotonic relationships between parameters and model outputs across a distribution of parameter sets. In general, the method computed partial correlations of the rank-transferred data [118].

LHS was used to generate *N*_*l*_ = 1, 000 evenly distributed parameter sets in which the locally influential parameters were varied and all other parameters were set to their nominal values given in either Tables 5 or 6. We defined the plausible range of each locally influential parameter to be 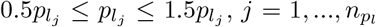, of its nominal value. For each of the *N*_*l*_ parameter sets, the system of ordinary differential equations was approximated numerically and each of the measurable outputs at each of the *n*_*t*_ days was stored.

We then computed a partial rank correlation coefficient, 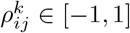, to quantify the monotonic association between the *k*-th output at time *t*_*i*_, *y*_*k*_(*t*_*i*_), *k* = 1, …, *n*_*y*_ and *i* = 1, …, *n*_*t*_, and the *j*-th locally influential parameter, 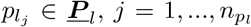. Values near 1 indicate a strong increasing monotonic relationship, whereas values near -1 indicate a strong decreasing monotonic relationship. Each partial rank correlation coefficient, 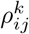, has an associated p-value, 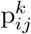, quantifying the significant of the correlation. Publicly available MATLAB functions were used to implement LHS-PRCC [119].

Globally influential parameters were stored in 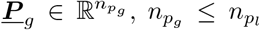, and are subset of the identified locally influential parameters, ***P***_*l*_. We identified an locally influential parameter 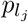 as globally influential by studying the frequencies of significant, strong monotonic relationships during the peak pollen season 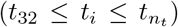. We defined the frequency of a significant, strong monotonic relationship of the measurable *k*-th output to parameter 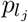 as

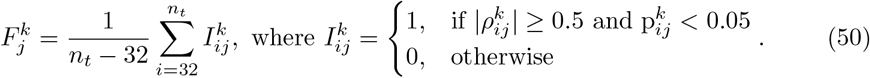

Specifically, if

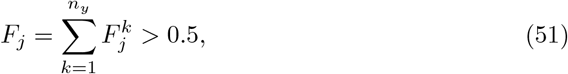

then the *j*-th locally influential parameter 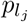 was identified as a globally influential parameter.

#### 3.2.2 Parameter Identifiability

Following parameter selection, we conducted structural identifiability analysis to assess whether the identified globally influential parameters can be uniquely determined from the available model outputs, independent of measurement noise or data limitations [120, 121]. Structural identifiability is a theoretical property of the model structure and provides insight into whether parameter estimation is feasible [121].

We used the open-sourced MATLAB toolbox GenSSI 2.0 to evaluate structural identifiability [120]. GenSSI 2.0 is an advancement of the toolbox GenSSI (Generating Series for testing Structural Identifiability) [121]. The toolbox couples the generating series approach with identifiability tableaus to determine whether parameters are either globally or locally identifiable under the assumed observation functions [120, 121].

#### 3.2.3 Parameter Fitting

In the final step of model calibration, we refined the globally-influential parameter values, ***P***_*g*_, to reproduce key features of the experimental data via parameter fitting. Again, we considered the experimental observed median, *M*^*k*^, first quartile, 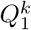, and third quartile, 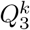, of the immune signal concentrations over the pollen season given in Table 7. These summary statistics capture both the central tendency and variability in the experimental data.

Model calibration was done via parameter fitting. We formulated our parameter fitting problem as a regression problem. For a given parameter set 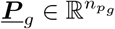, where all other parameters are set to their nominal values reported in Tables 5 and 6, we simulated our system of ordinary differential equations and stored the observable outputs at each day of the peak pollen season, *y*_*k*_(*t*_*i*_), *k* = 1, ‥, *n*_*y*_ and *t*_*i*_ = 32, …, *n*_*t*_. Each of the observable outputs was then normalized its respective experimental observed mean,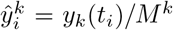. We then created the same summary statistics, 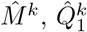, and 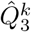, of the normalized simulated data using the MATLAB function quantile.

An objective function was built as the sum of squared differences between normalized simulated and experimental medians and quartiles,

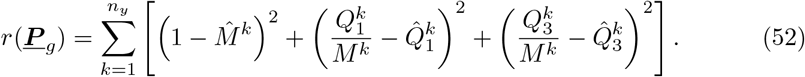

This objective function was minimized using a nonlinear least-squares approach implemented in MATLAB using the function lsqnonlin. To find the nominal best fit, we considered the admissible parameter set given by 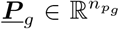, where 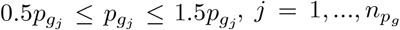, and 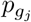 was set to its nominal value given in Tables 5 or 6. Differently, to find the the extreme best fit, we allowed the entries of <u>***P***</u>_*g*_ to range 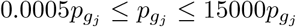.

## 4 Results

To begin, we present our model results using the nominal parameter values reported in Tables 5 and 6. We focus our discussion on the behavior of the experimentally observable immune signals 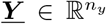, given in Equation (45). Then we report our model calibration findings.

### 4.1 Evolution of Measurable Immune Signals

Figure 4 shows the model predictions of experimentally measured inflammatory mediators, i.e., measurable immune signals ***Y*** (t). The eight different immune signal concentrations are plotted over the ragweed pollen season on a semi-logarithmic scale.

**Fig 4.**
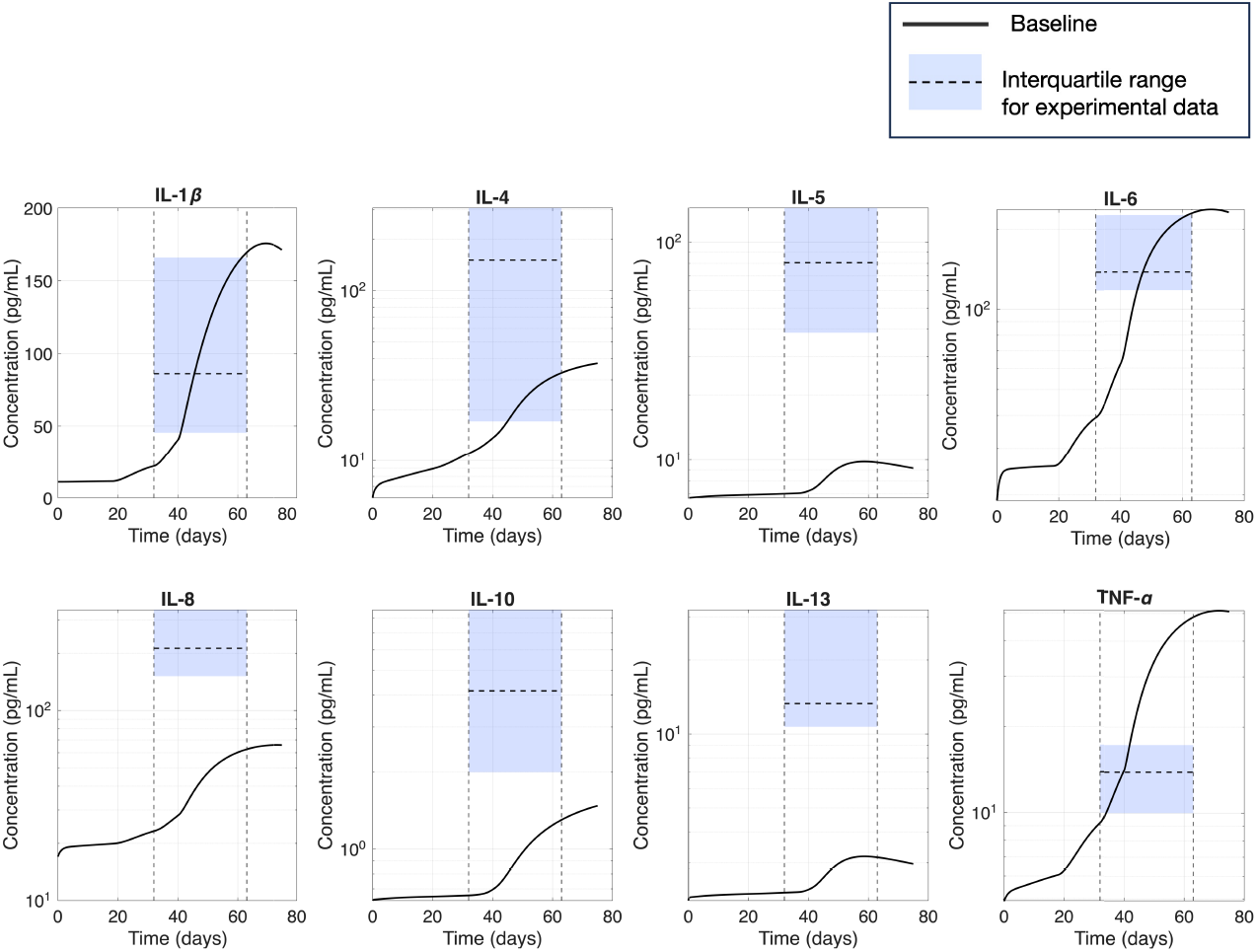
Evolution of the predicted observable immune signals, or inflammatory mediators, over the ragweed pollen season plotted on a semi-logarithmic scale. The horizontal dashed lines indicate the peak pollen season, (August 15th through September 15th, 32*≤ t ≤*63) [87]. The experimental data from Leonardi et al. [7] is represented by the shaded blue box plot with the median shown as the vertical dashed line.

The peak pollen season is indicated by the vertical dashed lines (August 15th (*t* = 32) through September 15th (*t* = 63)). Over the peak pollen season, we also show Leonardi et al. experimental data via a shaded blue box plot, where the median is denoted by the vertical dashed line [7]. The predicted concentrations of IL-1*β*, IL-4, IL-6, and TNF-*α* fall in the experimental ranges; whereas, the other predicted immune signal concentrations, specifically, IL-5, IL-8, IL-10, and IL-13, fall below the experimentally measured ranges.

Figure 5 illustrates the relative temporal evolution of each mediator over the ragweed pollen season via a spider plot. The simulated inflammatory mediators are expressed as a percentage of their respective seasonal maxima. Early in the pollen season (July 15th), the levels of immune signals remain near baseline indicating minimal inflammatory activation. As the pollen exposure increases, this leads to an increase in allergen concentration in the conjunctiva which is reflected by an increase in the immune signal percentage concentration in early-mid August. Peak inflammatory responses occur in early to mid September, corresponding to a peak in the pollen season as observed in Figure 2, followed by stabilization as allergen levels decline. In general, the expansion of the spider plot over time reflects increased activation across both pro-inflammatory and regulatory mediators consistent with a sustained, but regulated immune response.

**Fig 5.**
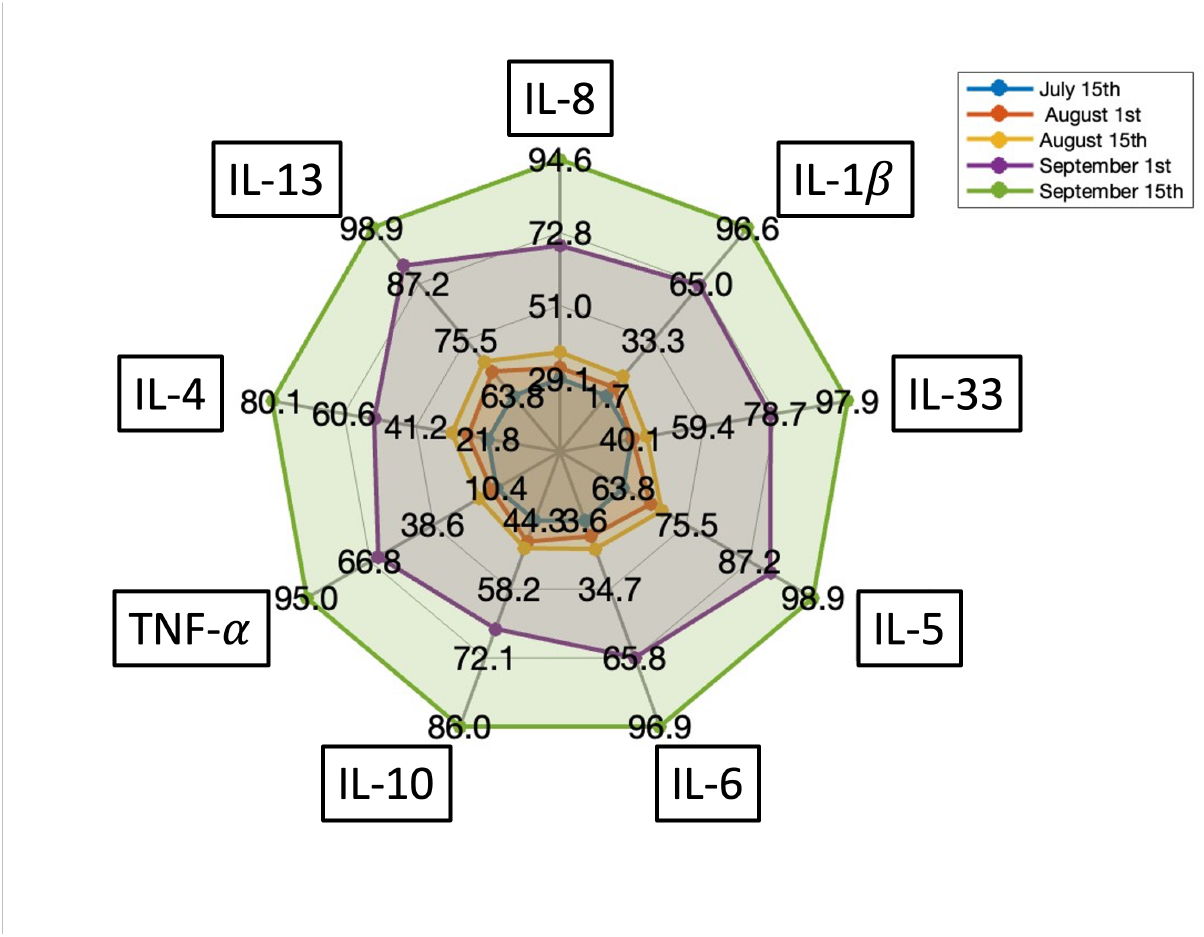
Evolution of the simulated inflammatory mediators, i.e., ***Y*** (*t*), during the ragweed pollen season. The color lines represent the percent increase relative to their maximum concentration at July 15th (blue line), August 1st (red line), August 15th (blue line), September 1st (purple line) and September 15th (green line).

Figure 6 compares model predictions to two different experimental data sets. In Figure 6**A**, we compare the box plots of the observable model predictions with Leonardi et al. data [7]. The simulated, or predicted box plots, were created from the daily immune signal concentration predictions over the peak ragweed pollen season. In general, the simulated inflammatory mediator concentrations fall within the observed experimental ranges. The median simulated levels of IL-1*β*, IL-4, and IL-6 lies within the inter-quartile range of the experimental data; deviations are observed for IL-5, IL-8, IL-10, IL-13, and TNF-*α*. The experimental measurements exhibit some differences which could be due to heterogeneity across patients.

**Fig 6.**
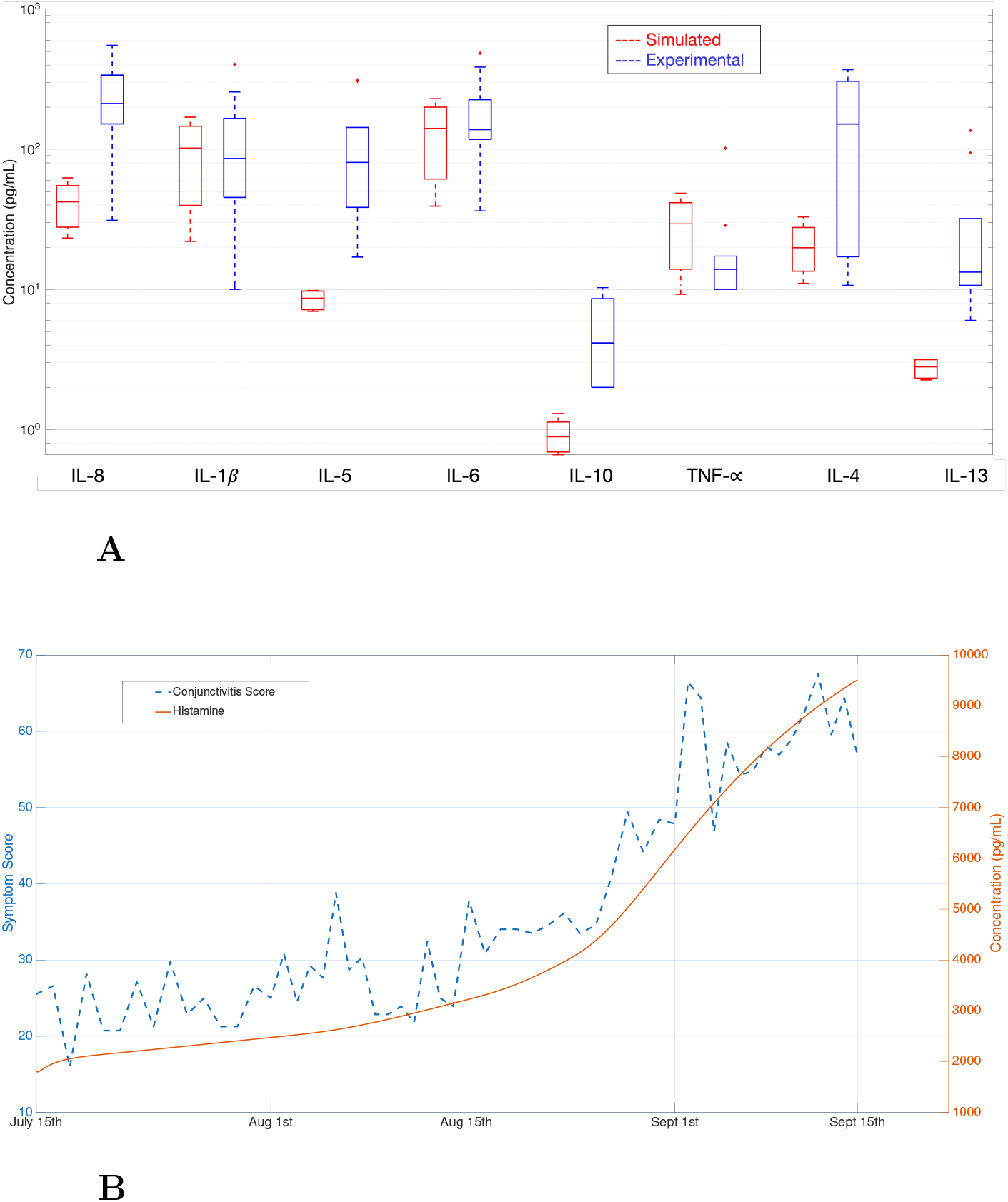
**(A)** Predicted immune signal concentrations for time points between August 15th (*t* = 32) to September 15th (*t* = 63), the peak ragwood pollen season (left red box plot) and experimental data shared by Leonardi et al. [7] (right blue box plot). **(B)** Comparison of the predicted histamine concentration levels with the conjunctivitis symptom score data (digitized from [87]). The Spearman’s rank correlation coefficient is *ρ* = 0.76.

Figure 6**B** compares the simulated histamine levels to daily conjunctivitis symptom scores collected in Bonini et al. [87]. Recall, the clinical manifestations captured by the symptom score are primarily mediated by histamine release [122]. While the symptom score displays greater day-to-day variability, both quantities increase during late August and early September and reach higher values near the peak of the pollen season. This similarity in overall trend is supported by a positive, significant Spearman rank correlation (*ρ* = 0.76, p *≤* 0.01).

### 4.2 Model Calibration

Figure 7 and Table 8 display the local sensitivity analysis findings. Figure 7 shows a bar chart of the sensitivity index, 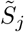, for each model parameter, *p*_*j*_. The parameters are in ascending order of sensitivity index with most locally sensitive parameters listed first. The parameter list is given in Table 8 along with the numerical sensitivity index value. Among the *n*_*p*_ = 77 parameters perturbed, only 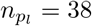 parameters out of the *n*_*p*_ = 77 are locally sensitive to the model outputs. The majority of these influential parameters correspond to production or depletion rates of the inflammatory mediators, as summarized in Table 8.

**Fig 7.**
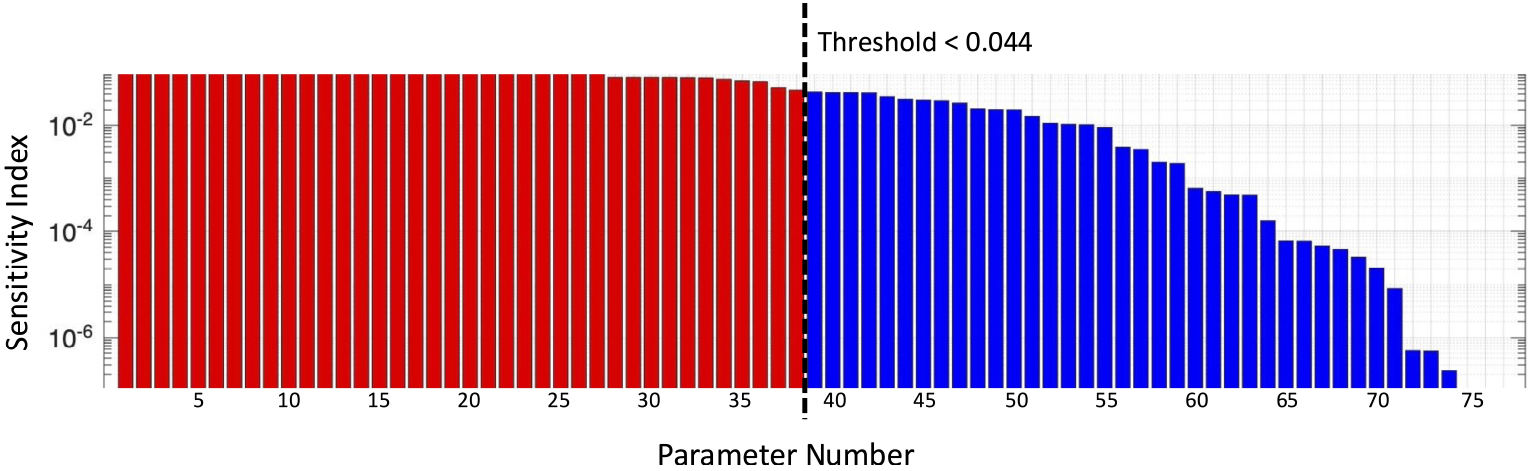
Bar plot of the sensitivity index value 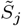 for each parameter *p*_*j*_. The parameters are ranked in decreasing order of their sensitivity index value and listed in Table 8. The parameters identified as locally influential, Equation (49), are highlighted in red.

Our global sensitivity analysis results are shown in Figure 8 and reported in Table 9. Figure 8**A** plots the temporal evolution of PRCC values of the measurable output IL-5 to each of the locally influential parameters over the pollen season (i.e., 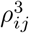, for *i* = 1, …, *n*_*t*_ and *j* = 1, …, 38). The three-dimensional plot illustrates how the strength and direction of output-parameter correlations change over time. Figure 8**B** summarizes the frequency of significant, strong correlations of each outputs to each locally influential parameter. Each bar corresponds to a different locally influential parameter. The different colors within one bar are the frequencies of the different outputs. Parameters 1, 2, 5, 6, 12, 15, 17, 19–21, 23, 24, 26, and 33–35 were identified as globally sensitive as summarized in Table 9 (complete results in Table B1); that is,

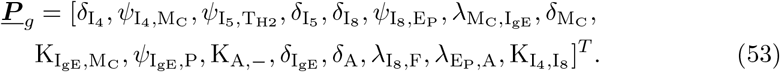

**Table 9.**
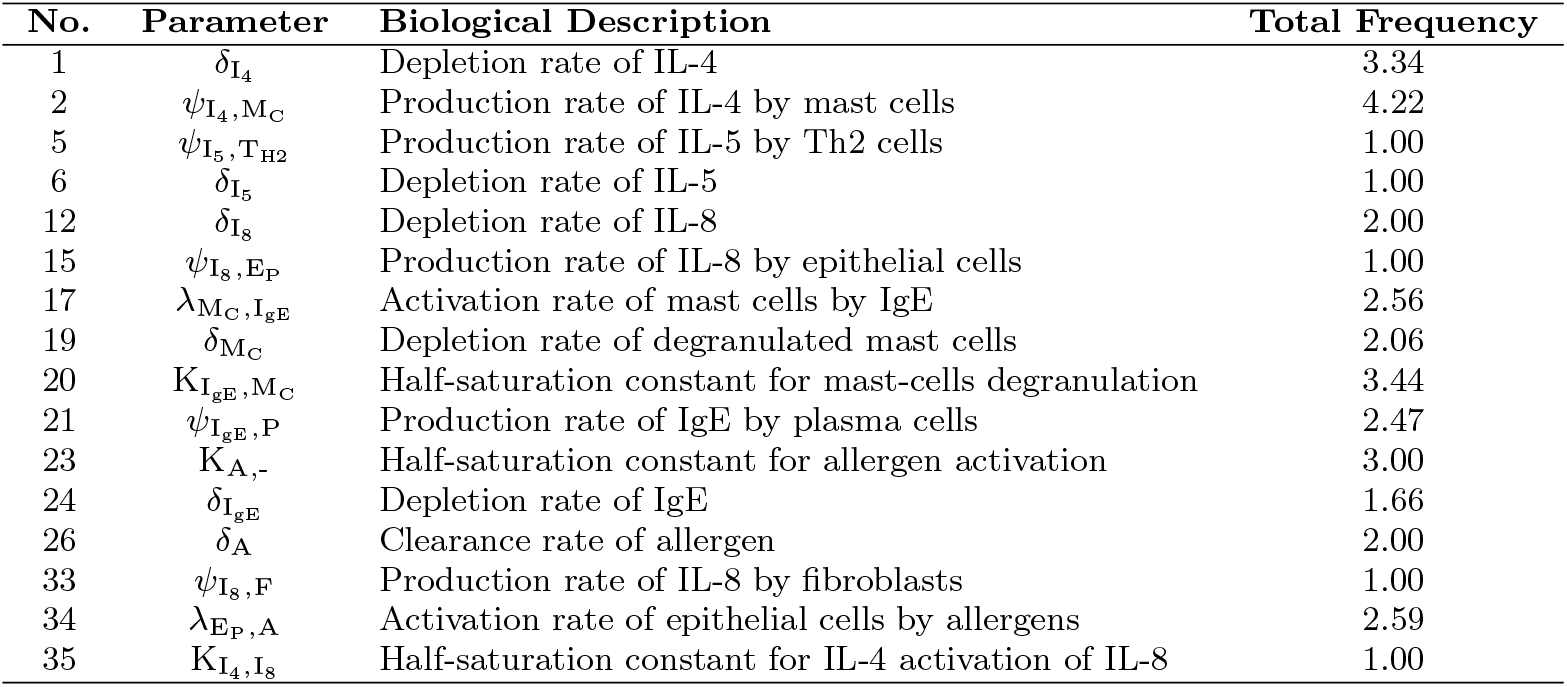
Globally influential parameters identified by LHS-PRCC analysis. For each of these locally influential parameters, the total frequency of significant, strong correlations over the pollen season is greater than 0.5 (i.e., *F*_*j*_ *>* 0.5, defined in Equation (51)).

| No. | Parameter | Biological Description | Total Frequency |
| --- | --- | --- | --- |
| 1 | $\delta_{I_4}$ | Depletion rate of IL-4 | 3.34 |
| 2 | $\psi_{I_4, M_C}$ | Production rate of IL-4 by mast cells | 4.22 |
| 5 | $\psi_{I_5, T_{H2}}$ | Production rate of IL-5 by Th2 cells | 1.00 |
| 6 | $\delta_{I_5}$ | Depletion rate of IL-5 | 1.00 |
| 12 | $\delta_{I_8}$ | Depletion rate of IL-8 | 2.00 |
| 15 | $\psi_{I_8, E_P}$ | Production rate of IL-8 by epithelial cells | 1.00 |
| 17 | $\lambda_{M_C, I_{gE}}$ | Activation rate of mast cells by IgE | 2.56 |
| 19 | $\delta_{M_C}$ | Depletion rate of degranulated mast cells | 2.06 |
| 20 | $K_{I_{gE}, M_C}$ | Half-saturation constant for mast-cells degranulation | 3.44 |
| 21 | $\psi_{I_{gE}, P}$ | Production rate of IgE by plasma cells | 2.47 |
| 23 | $K_{A, -}$ | Half-saturation constant for allergen activation | 3.00 |
| 24 | $\delta_{I_{gE}}$ | Depletion rate of IgE | 1.66 |
| 26 | $\delta_A$ | Clearance rate of allergen | 2.00 |
| 33 | $\psi_{I_8, F}$ | Production rate of IL-8 by fibroblasts | 1.00 |
| 34 | $\lambda_{E_P, A}$ | Activation rate of epithelial cells by allergens | 2.59 |
| 35 | $K_{I_4, I_8}$ | Half-saturation constant for IL-4 activation of IL-8 | 1.00 |

**Fig 8.**
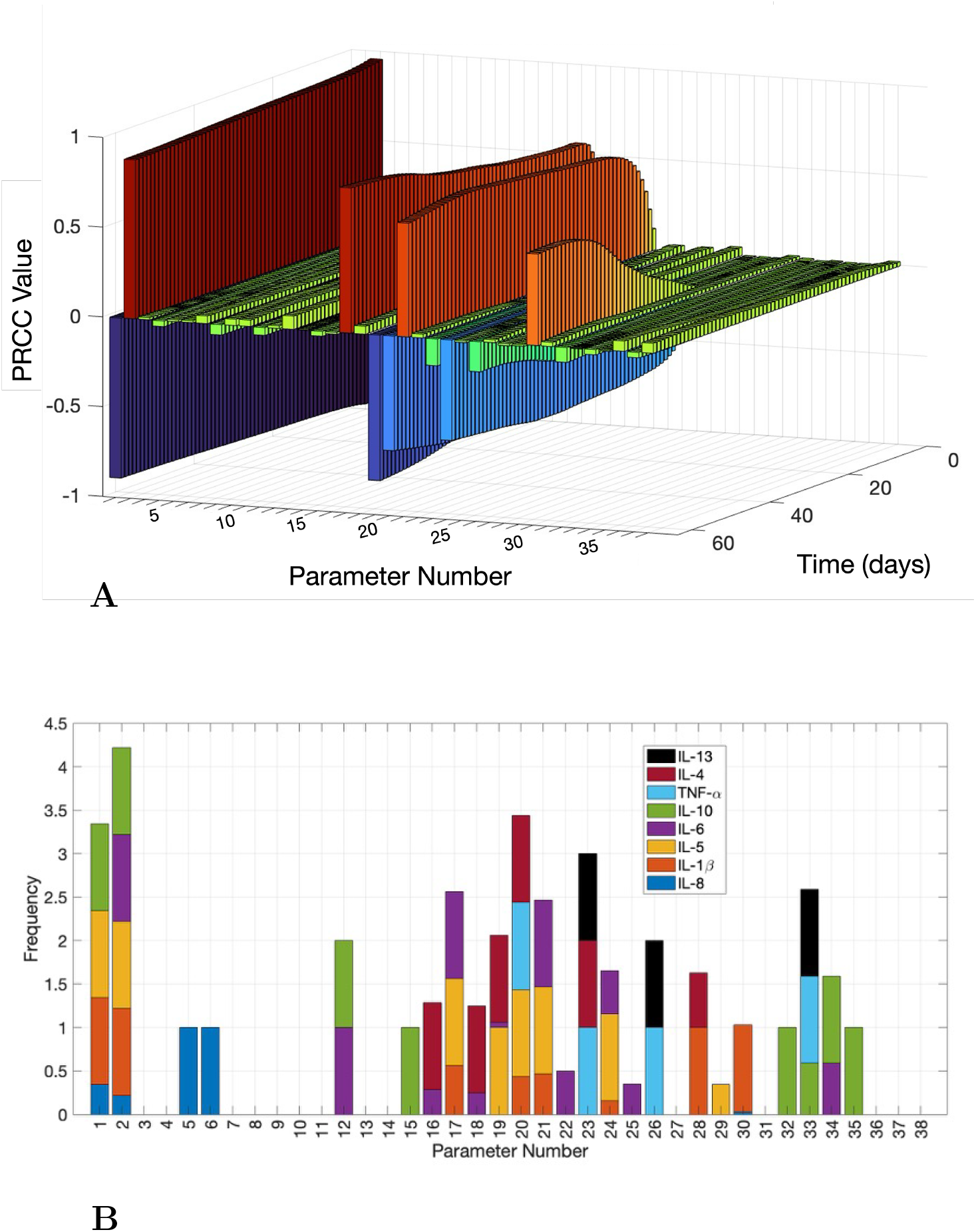
**(A)** The PRCC of IL-5 to each locally influential parameter over the duration of the pollen season (i.e., 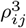, for *i* = 1, …, *n*_*t*_ and *j* = 1, …, 38). Recall, 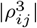*>* 0.5 indicates a strong influence of the *j*-th parameter on model output IL-5 at the *i*-th day. **(B)** The frequency of a significant, strong correlation of the *k*-th output to the *j*-th locally influential parameter over the peak pollen season (i.e., 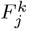, for *j* = 1, …, 38 and *k* = 1, …, 8, defined in Equation (50)). Each bar corresponds to a different locally influential parameter. The different colors in a bar denote the frequencies of different outputs.

Next, we report the structural indentifiability of the globally influential parameters, 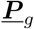. The structural identifiability calculation required fifth-order Lie derivatives to assess parameter identifiability (see the structural identifiability tableau shown in Figure B1). We find that the model is at least locally identifiable for each of the globally influential parameters. Parameters 5, 6, and 23 (production and death rates of IL-5 and the half saturation constant for allergen activation) are identified as globally identifiable. With each parameter at locally identifiable, we moved forward with model calibration.

Figure 9 summarizes the results of the model calibration in the form of a forest plot, illustrating the ranges of the calibrated globally influential parameters relative to their baseline values. For each parameter, the nominal best fit estimate and the extreme best fit value are shown as a percentage of the baseline parameter value. The shaded region indicates the range of parameter values that represent +*/-* 50% of the baseline values (also referred to as nominal values) which aligns with biological relevant ranges.

**Fig 9.**
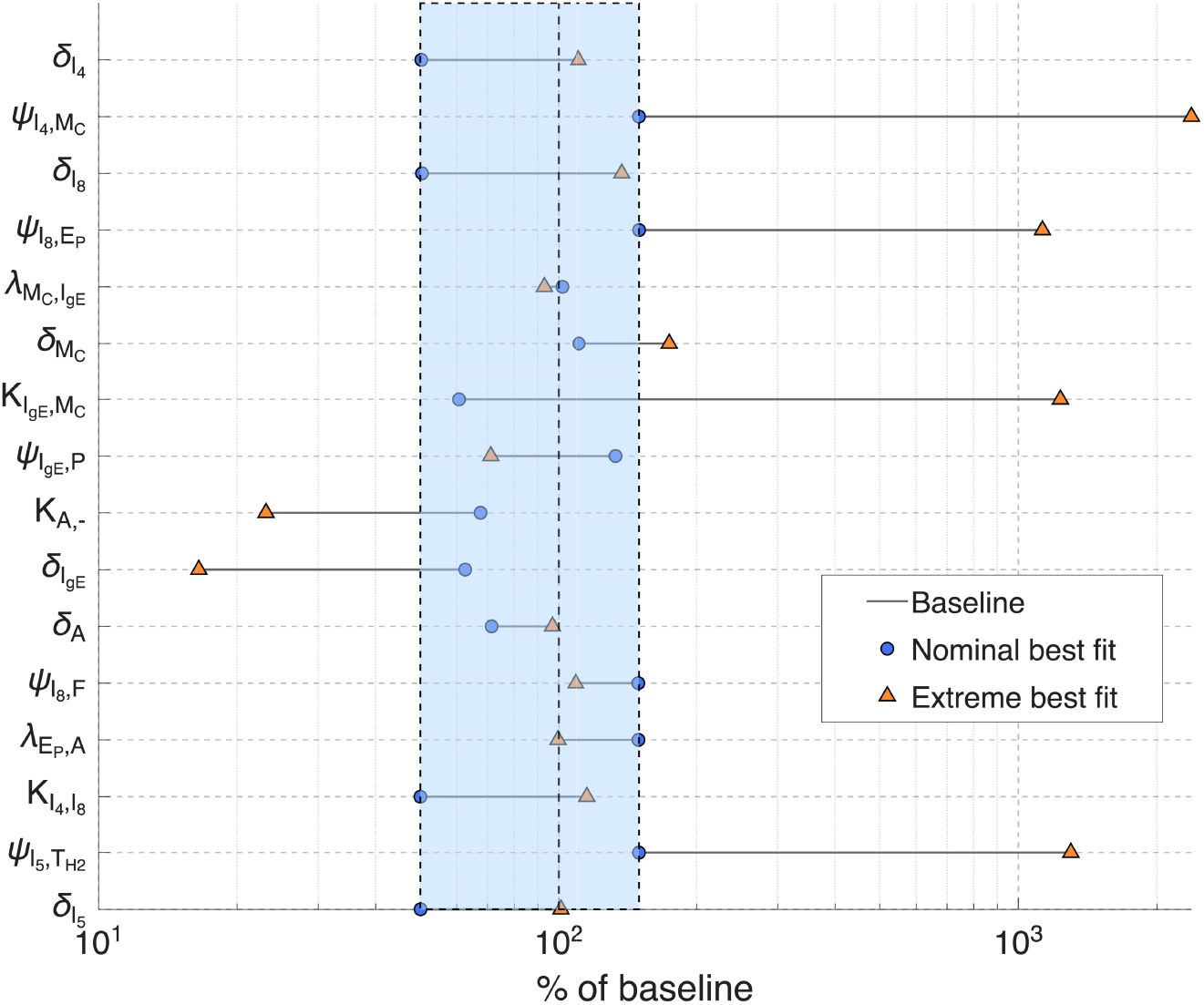
Forest plot of calibrated model parameters shown as a percentage of their baseline values. Blue circles denote nominal best-fit parameter estimates, while orange triangles indicate extreme best-fit values obtained during calibration. Horizontal lines represent the range of parameter values. The shaded region highlights parameter ranges of 50% to 150% of the nominal value.

**Fig 10.**
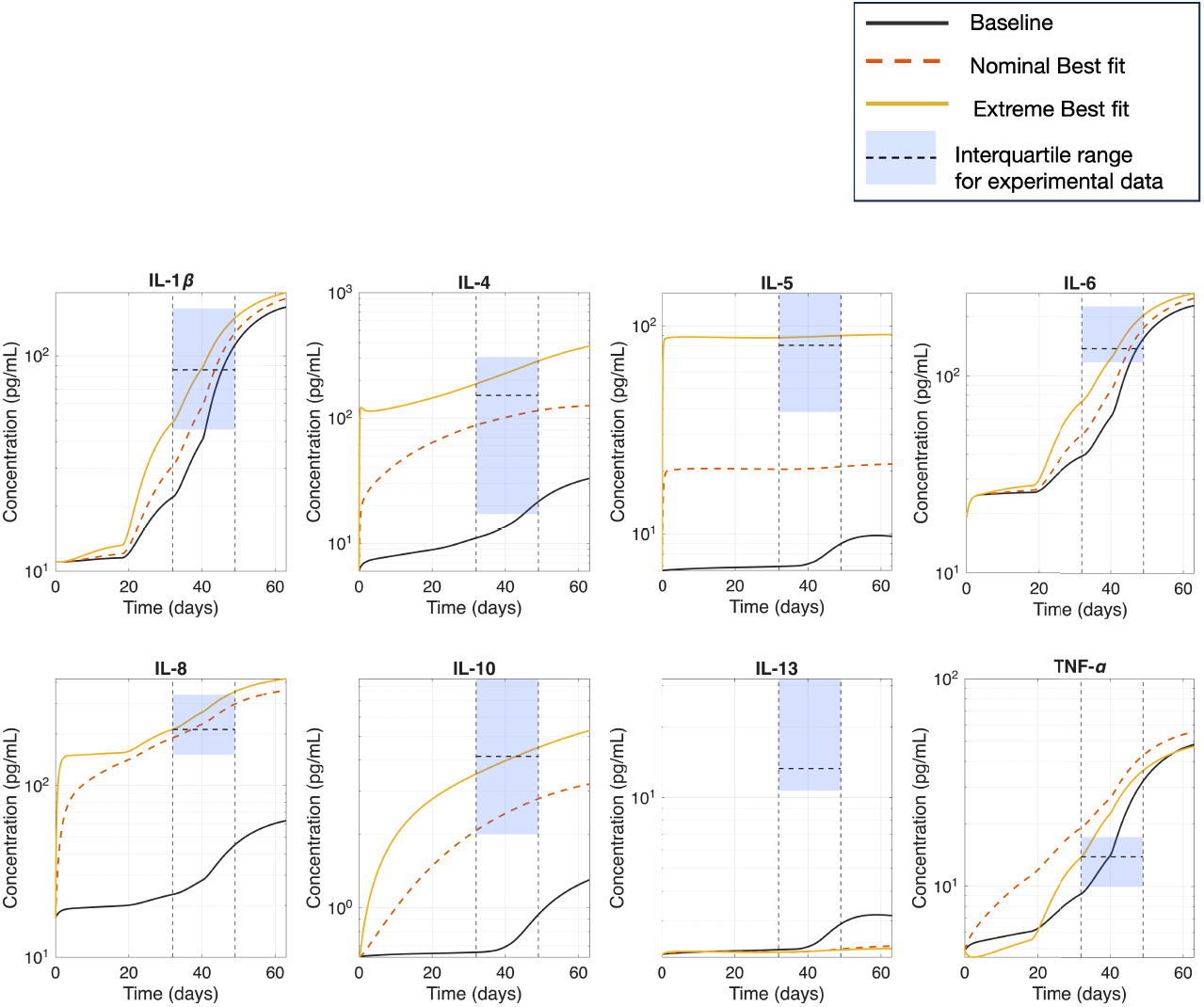
Progression of the simulated key inflammatory mediators concentrations over the pollen season with calibrated parameter values. Model predictions are shown for the baseline parameter set (black solid line), the nominal-best-fit parameter set (red dotted line), and the extreme-best-fit parameter set (yellow solid line). Experimental measurements are summarized by their interquartile ranges.

Overall, the majority of calibrated parameters remain within one order of magnitude of their baseline values, indicating that a nominal best fit estimate parameter adjustments are able to capture the model outputs with experimental data. However, a subset of parameters, specifically, 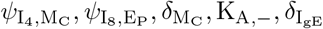, and 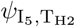, exhibits a wider range suggesting reduced constraint by the available data within the model structure. These results highlight parameter-specific differences in calibration sensitivity and underscore the importance of accounting for uncertainty when interpreting fitted parameter values.

Figure 11 compares experimental measurements with simulated concentrations obtained using baseline, nominal-best-fit, and extreme-best-fit parameter sets. For IL-5, IL-8, and IL-10, both the nominal- and extreme-best-fit calibrations shift the median simulated concentrations closer to the experimental distributions indicating improved agreement relative to the baseline model. In these cases, calibration primarily affects the central tendency of the simulated outputs while maintaining variability within a biological range. In contrast, calibration does not improve the ranges for the simulated IL-13. The calibrated simulations deviate further from the corresponding experimental distribution suggesting that the parameters adjusted during calibration are insufficient to capture the observed IL-13 dynamics.

**Fig 11.**
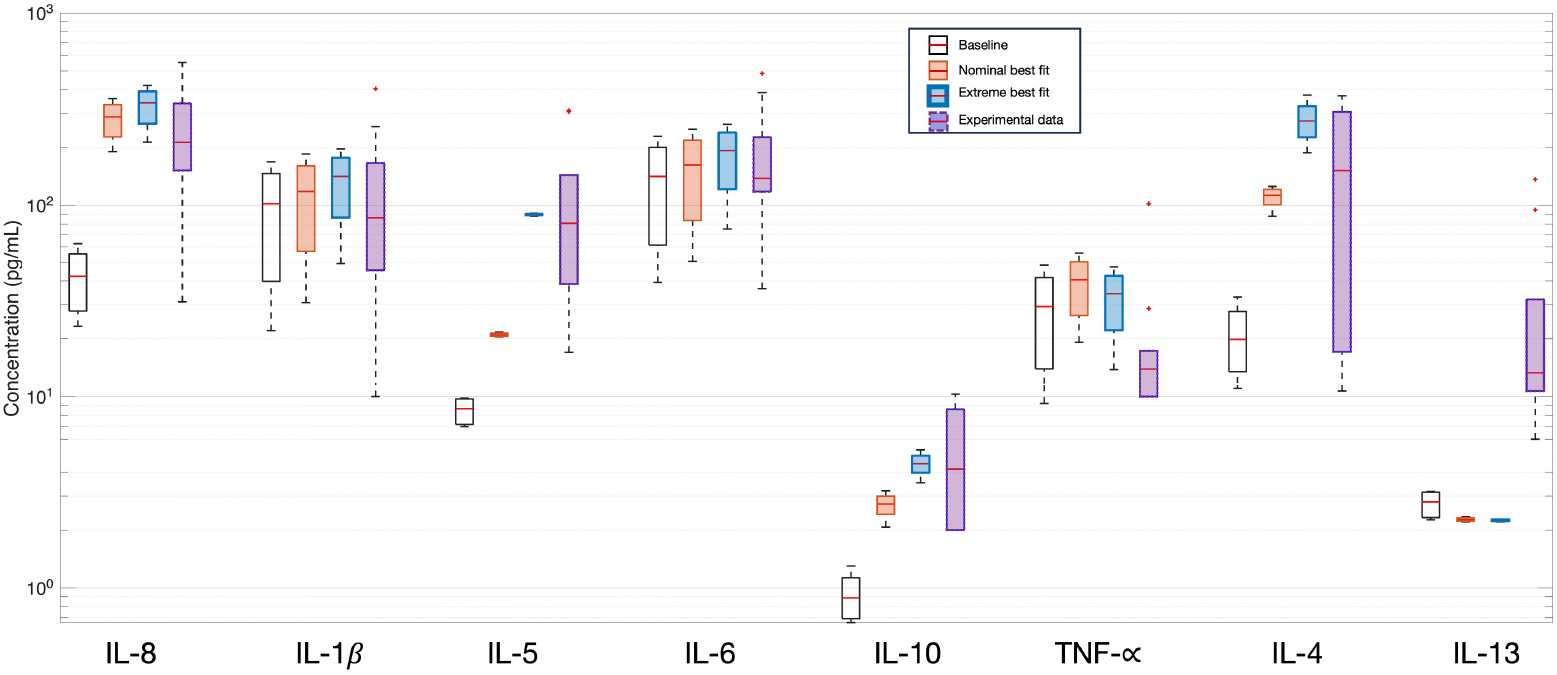
Comparison of simulated and experimental concentrations following model calibration. Box plots show experimental measurements alongside simulated concentrations obtained using baseline parameters (clear box plot), nominal-best-fit parameters (orange box plot), extreme-best-fit parameters (blue bold box plot) and experimental data (purple dotted box plot) for the key inflammatory mediators.

## 5 Discussion and Conclusions

In this work, we developed a mechanistic model to describe the immune response of the conjunctiva during SAC. By integrating allergen exposure dynamics with immune cell activation, immune signaling, and regulatory feedback mechanisms, the model provides a quantitative framework for examining how inflammatory processes evolve over the course of a pollen season. To our knowledge, this is one of the first system-level mathematical models of SAC.

Model simulations reproduce key qualitative features of SAC progression, including minimal inflammatory activity early in the pollen season, followed by a coordinated increase in pro-inflammatory mediators as allergen exposure intensifies. The temporal evolution of immune signals, such as IL-4, IL-5, IL-6, IL-8, and TNF-*α*, reflects the underlying Th2-driven immune response characteristic of allergic conjunctivitis. We find the inflammatory activity in the conjunctiva is due to a sustained immune activation that is modulated, but not fully suppressed, by regulatory immune signals such as IL-10 and TGF− *β*.

We found that conjunctivitis symptom score data was strongly correlated with our predicted histamine levels (see Figure 6**B**). The conjuctivitis symptom score is an indirect measurement of the inflammatory burden over time. A strong correlation suggests that the model can capture clinically relevant trends in SAC progression.

Our sensitivity analysis found a subset of parameters, primarily activation and production rates of immune signals, are influential on the observable immune signals. This is consistent with immunological expectations as immune signals amplifies the immune response through nonlinear feedback mechanisms [1, 3]. We found that IL-4 depletion rate and its production rate via mast cell degranulation had the strongest influence on the observable immune signals (see Table 9). Recall, in our model, IL-4 is the key driver of inflammation through the polarization of Th2 cells (pathway 14), the activation of B cells (pathway 37), and differentiation of plasma cells (pathway 30) which all contribute to the production of IgE (pathway 38).

The sensitivity of parameters associated with the depletion and production of IL-4 is consistent with the clinical landscape. Dupilumab, a monoclonal antibody targeting the shared IL-4R*α* subunit of the IL-4 and IL-13 receptor complex, has demonstrated great efficacy in allergic inflammatory diseases including moderate-to-severe asthma, and atopic dermatitis [123]. This further reinforce our model’s identification of IL-4 signaling as a potential therapeutic target.

Similarly, we found the production rates of IL-5 and IL-8 are also globally influential to the observable immune signals. In our SAC model, IL-5 is a principal driver for eosinophils activation (pathway 28) meaning that the sensitivity of IL-5 impacts the model capacity to regulate eosinophil-mediated tissue damage. This finding aligns with the clinical efficacy of mepolizumab (anti-IL5R*α*) which is a receptor blocker for the IL-5 receptor [124]. These are established treatments for severe eosinophilic associated disorder and asthma [124].

The global influence of both IL-4 and IL-5 suggests the inflammatory dynamics modeled are not governed by a single dominant pathway but rather emerge from a distributed network of sensitive nodes. Taken together, the consistency between the model’s sensitivity landscape and the established clinical targets of allergic inflammatory disease (asthma and eosinophilic disorders) support our model’s underlying biology. This further implies that therapeutic strategies targeting only one mediator may only achieve incomplete suppression in limiting inflammatory activity for SAC, and that a combination of interventions may be necessary to achieve effective suppression of the overall inflammatory activity for the disease

We found, for several immune signals, model calibration successfully shifts model predictions toward experimental distributions. However, residual discrepancies, particularly for IL-5 and IL-13, highlight the biological complexity and suggest that additional mechanisms may play a role in shaping these responses. In particular, it is known that mast cells and eosinophils produce both IL-5 and IL-13 [1]. Additionally, we note that differences between the model predictions and the experimental data could be influenced by limitations of the available experimental dataset. Limitations include a small cohort and lacks detailed patient-specific covariates such as disease severity, age, and prior allergic history.

We made several simplifying assumptions. For example, we treated the conjunctival environment as a well-mixed compartment and therefore spatial heterogeneity was not explicitly modeled. We also made assumptions about the immune homeostatic state. While these assumptions are common in mechanistic immune modeling, they may influence quantitative predictions and should be revisited as additional data become available.

Future extensions of our modeling framework could incorporate spatial heterogeneity within the conjunctiva microenvironment or patient-specific variability to better capture localized immune responses at the ocular surface. Additionally, the model could be extended to investigate therapeutic interventions, such as antihistamines, mast-cell stabilizers, or biological targeting certain immune signals, enabling in silico evaluation of treatment timing and efficacy. Such a study focused on antihistamines was done by Gunputh [125].

## Acknowledgments

We thank Dr. Andrea Leonardi for sharing the data published in [7]. N.G. acknowledges the financial support of the RIT Steven M. Wear Endowed Graduate Fellowship.

## Appendix A Parameter Estimation

### A.1 Allergen-driven immune cell change-of-state rates, 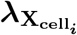,A

The homeostatic immune system response describes the dynamics when no allergens are presents. Consequently, allergen-activated immune cell rates terms will be zero and we cannot solve for 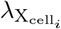,A. Therefore, we assume these activation rates are equal to the immune cell death rates. The specific rates determine this way are *λ*_D,A_ (pathway 2, parameter number 47), 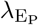,A (pathway 1, parameter number 34), and 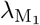, A (pathway 3, parameter number 13).

### A.2 Immune-signal driven immune cell change-of-state rates, 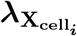, 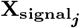

We assume the homeostatic immune signals are different from zero; that is, 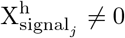 for *j ≠* 1. Consequently, at homeostasis, the immune signal activation function is different from zero; i.e., 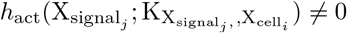 (see Equation (5)). Therefore, we determine the immune-signal driven immune cell change-of-state rates, 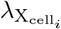, 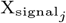, by setting 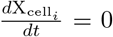 and ***X*** = ***X***^h^, and then solving for 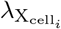, 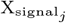. The specific rates determined this way are 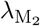, I_4_ (pathway 40, parameter number 45) and 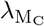, 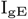 (pathway 32, parameter number 17).

For the Th2 cell population dynamics, Equation (30) contains four different immune signal activation rates, 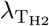, 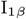 (pathway 12, parameter number 51), 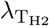, I_4_ (pathway 14, parameter number 39), 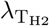, I_6_ (pathway 10, parameter number 36), 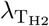, I_33_ (pathway 11, parameter number 76). It is argued that IL-4 and IL-6 are the major driver of the Th2 polarization/activation, while IL-1*β* and IL-33 are the secondary/minor contributors in the activation of Th2 cells [1, 3]. Thus, we assume that IL-6 contributes to 70% activation of Th2 cells, IL-4 contributes to 20%, and both IL-33 and IL-1*β* contributes 5%. Using these weights, we solve for the four rates at homeostasis as described above.

### A.3 Inhibitory effect of IL-10, 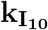, −

IL-10 can limit Th2 responses by negatively regulating Th2 clones and reducing the secretion of IL-4 and IL-5 in allergic inflammation models [1, 32, 33]. In ocular allergy, IL-10 contributes to the control of Th2-mediated inflammation, yet tear measurements show that IL-10 levels are insufficient to suppress IL-4 production completely [5, 40]. IL-10 is a well-established suppressor of TNF-*α* production in monocytes and macrophages, acting through inhibition of NF-*κ*B signaling and transcriptional repression of pro-inflammatory genes [6, 126]. IL-10 also reduces TNF-*α* release from IgE-activated human mast cells [41]. Clinical ocular allergy studies report a lower IL-10/TNF-*α* ratio in tears, indicating that IL-10 signaling is insufficient to fully counterbalance TNF-*α*–driven inflammation [5, 40].

Borish et al. has shown that IL-10 exhibits weak inhibitory activity at a concentration of 10 pg/mL [127]. Therefore, we assume IL-10 will similarly exert the same weak inhibition on (i) mast cell degranulation, (ii) TNF-*α* production and (iii) IL-4 production. To represent the suppressive effect of IL-10, we set the inhibition function to approximately 10% at IL-10 concentration of 10 pg/mL to reflect a weak inhibition of, and solve for the inhibition parameter 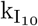, *−* as follows:

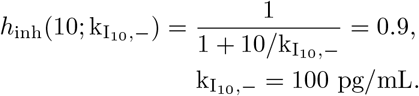

Consequently, 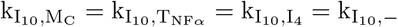.

### A.4 Inhibitory effect of TGF-*β*, 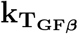,**T**_**H2**_

Kunzmann et al. showed that TGF-*β*1 exerts a strong, dose-dependent suppression of Th2 responses in human CD4^+^ T cells, and that this suppression is further enhanced by histamine signaling through the H_2_ receptor [27]. In their study, histamine dose-dependently increased TGF-*β*1 responsiveness [27]. These data support the view that TGF-*β* can strongly suppress Th2 differentiation when present at sufficient concentrations. Gorelik also showed in their study that TGF *− β* causes a near complete inhibition of Th2 cells differentiation in the CD4+T cell culture [128].

To capture this behavior, we assume that 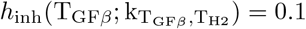 allows for up to approximately 90% suppression of the Th2 activation term. Consistent with studies showing TGF-*β* at 1–10 ng/mL (1,000–10,000 pg/mL) suppresses Th2 activation, we used a concentration of T_GF*β*_ = 4, 000 pg/mL to solve for 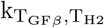. Thus, the choice of 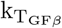, T_H2_ is made so that the model can reproduce the strong (near-order-of-magnitude) suppression of Th2 responses reported for TGF-*β*1 in vitro, while still allowing residual Th2 activity under conditions of high TGF-*β*.

### A.5 Immune signal production rates, 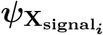, 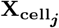

We assume the homeostatic immune cells and signals are different from zero; that is, 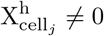 and 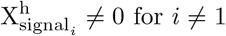. Therefore, we determine the immune signal production rates, 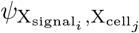, by setting 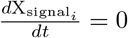 and ***X*** = ***X***^h^, and then solving for 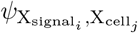. The specific rates determined this way are 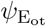, F (pathway 24, parameter number 66), *ψ*_H,MC_ (pathway 36, parameter number 72), 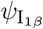, M_1_ (pathway 29, parameter number 4), 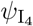, M_C_ (pathway 23, parameter number 2), 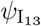, T_H2_ (pathway 19, parameter number 10), 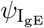, P (pathway 38, parameter number 21), and 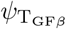, M_2_ (pathway 39, parameter number 29).

In the IL-6 immune signal dynamics (see Equation (37)), there are two production rates, 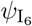, D (pathway 6, parameter number 27) and 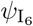, M_1_ (pathway 7, parameter number 16). We assume the rates are same value and solve for that value at homeostasis.

The IL-8 immune signal dynamics (see Equation (38)) has two production rates, 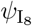, F (pathway 25 and parameter number 33) and 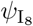, E_P_ (pathway 9 and parameter 40 number 15). Cubitt et al. reported that human corneal keratocytes (stromal fibroblast-like cells) produced approximately 33 times more IL-8 than corneal epithelial cells under comparable activation conditions [129]. We assume a similar relationship by setting 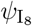, 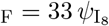, E_P_. Then we solve for the rates at homeostasis.

In the IL-10 immune signal dynamics (see Equation (39)), there are two production rates, 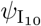, T_H2_ (pathway 16 and parameter number 14) and 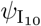, M_2_ (pathway 17, parameter number 38). We assume the rates are the same value and solve for that value at homeostasis.

Similarly, in the TNF-*α* immune signal dynamics (see Equation (44)), there are two production rate, 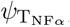, M_1_ (pathway 33, parameter number 18) and 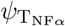, M_C_ (pathway 34, parameter number 22). We assume the rates are the same value and solve for that value at homeostasis.

## Appendix B Additional Results

### B.1 Global Sensitivity Analysis

Table B1 displays the computed fequency of a strong, significant correlation between each immune signals and each locally sensitive parameter.

### B.2 Structural Identifiability

Figure B1 is the identifiability tableau. A black line at location (*i, j*) means the non-zero generating series coefficient *i* depends on parameter *j*. The Jacobian was full rank after computing the fifth-order Lie derivatives.

**Table B1.**
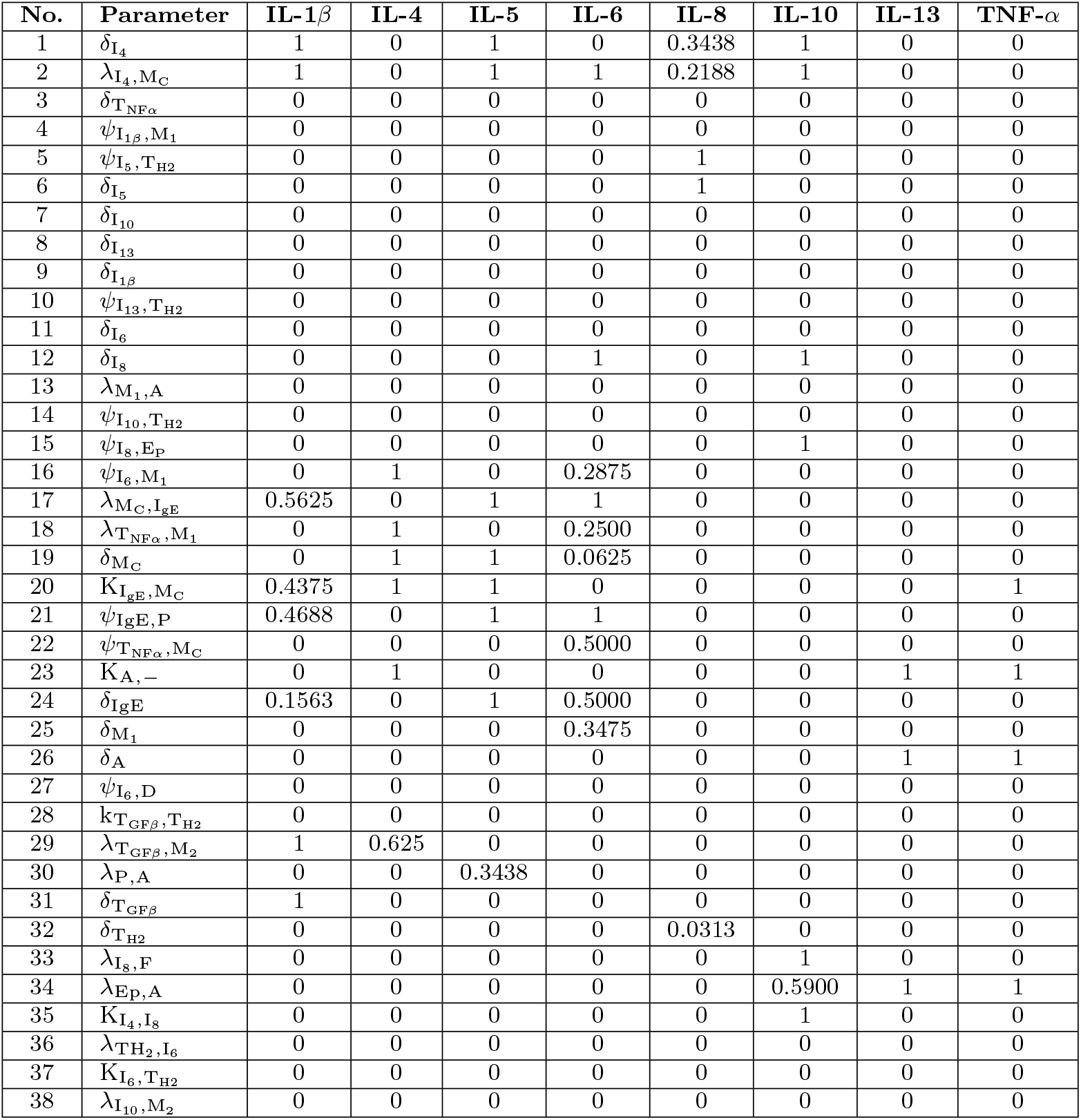
The computed frequency of strong, significant correlations between each observable output and each locally influential parameter during the time period *t* = 32 to *t* = 63 days.

| No. | Parameter | IL-1β | IL-4 | IL-5 | IL-6 | IL-8 | IL-10 | IL-13 | TNF-α |
| --- | --- | --- | --- | --- | --- | --- | --- | --- | --- |
| 1 | $\delta_{I_4}$ | 1 | 0 | 1 | 0 | 0.3438 | 1 | 0 | 0 |
| 2 | $\lambda_{I_4, M_C}$ | 1 | 0 | 1 | 1 | 0.2188 | 1 | 0 | 0 |
| 3 | $\delta_{T_{NF\alpha}}$ | 0 | 0 | 0 | 0 | 0 | 0 | 0 | 0 |
| 4 | $\psi_{I_{1\beta}, M_1}$ | 0 | 0 | 0 | 0 | 0 | 0 | 0 | 0 |
| 5 | $\psi_{I_5, T_{H2}}$ | 0 | 0 | 0 | 0 | 1 | 0 | 0 | 0 |
| 6 | $\delta_{I_5}$ | 0 | 0 | 0 | 0 | 1 | 0 | 0 | 0 |
| 7 | $\delta_{I_{10}}$ | 0 | 0 | 0 | 0 | 0 | 0 | 0 | 0 |
| 8 | $\delta_{I_{13}}$ | 0 | 0 | 0 | 0 | 0 | 0 | 0 | 0 |
| 9 | $\delta_{I_{1\beta}}$ | 0 | 0 | 0 | 0 | 0 | 0 | 0 | 0 |
| 10 | $\psi_{I_{13}, T_{H2}}$ | 0 | 0 | 0 | 0 | 0 | 0 | 0 | 0 |
| 11 | $\delta_{I_6}$ | 0 | 0 | 0 | 0 | 0 | 0 | 0 | 0 |
| 12 | $\delta_{I_8}$ | 0 | 0 | 0 | 1 | 0 | 1 | 0 | 0 |
| 13 | $\lambda_{M_1, A}$ | 0 | 0 | 0 | 0 | 0 | 0 | 0 | 0 |
| 14 | $\psi_{I_{10}, T_{H2}}$ | 0 | 0 | 0 | 0 | 0 | 0 | 0 | 0 |
| 15 | $\psi_{I_8, E_P}$ | 0 | 0 | 0 | 0 | 0 | 1 | 0 | 0 |
| 16 | $\psi_{I_6, M_1}$ | 0 | 1 | 0 | 0.2875 | 0 | 0 | 0 | 0 |
| 17 | $\lambda_{M_C, I_{gE}}$ | 0.5625 | 0 | 1 | 1 | 0 | 0 | 0 | 0 |
| 18 | $\lambda_{T_{NF\alpha}, M_1}$ | 0 | 1 | 0 | 0.2500 | 0 | 0 | 0 | 0 |
| 19 | $\delta_{M_C}$ | 0 | 1 | 1 | 0.0625 | 0 | 0 | 0 | 0 |
| 20 | $K_{I_{gE}, M_C}$ | 0.4375 | 1 | 1 | 0 | 0 | 0 | 0 | 1 |
| 21 | $\psi_{I_{gE}, P}$ | 0.4688 | 0 | 1 | 1 | 0 | 0 | 0 | 0 |
| 22 | $\psi_{T_{NF\alpha}, M_C}$ | 0 | 0 | 0 | 0.5000 | 0 | 0 | 0 | 0 |
| 23 | $K_{A, -}$ | 0 | 1 | 0 | 0 | 0 | 0 | 1 | 1 |
| 24 | $\delta_{I_{gE}}$ | 0.1563 | 0 | 1 | 0.5000 | 0 | 0 | 0 | 0 |
| 25 | $\delta_{M_1}$ | 0 | 0 | 0 | 0.3475 | 0 | 0 | 0 | 0 |
| 26 | $\delta_A$ | 0 | 0 | 0 | 0 | 0 | 0 | 1 | 1 |
| 27 | $\psi_{I_6, D}$ | 0 | 0 | 0 | 0 | 0 | 0 | 0 | 0 |
| 28 | $k_{T_{GF\beta}, T_{H2}}$ | 0 | 0 | 0 | 0 | 0 | 0 | 0 | 0 |
| 29 | $\lambda_{T_{GF\beta}, M_2}$ | 1 | 0.625 | 0 | 0 | 0 | 0 | 0 | 0 |
| 30 | $\lambda_{P, A}$ | 0 | 0 | 0.3438 | 0 | 0 | 0 | 0 | 0 |
| 31 | $\delta_{T_{GF\beta}}$ | 1 | 0 | 0 | 0 | 0 | 0 | 0 | 0 |
| 32 | $\delta_{T_{H2}}$ | 0 | 0 | 0 | 0 | 0.0313 | 0 | 0 | 0 |
| 33 | $\lambda_{I_8, F}$ | 0 | 0 | 0 | 0 | 0 | 1 | 0 | 0 |
| 34 | $\lambda_{E_P, A}$ | 0 | 0 | 0 | 0 | 0 | 0.5900 | 1 | 1 |
| 35 | $K_{I_4, I_8}$ | 0 | 0 | 0 | 0 | 0 | 1 | 0 | 0 |
| 36 | $\lambda_{T_{H2}, I_6}$ | 0 | 0 | 0 | 0 | 0 | 0 | 0 | 0 |
| 37 | $K_{I_6, T_{H2}}$ | 0 | 0 | 0 | 0 | 0 | 0 | 0 | 0 |
| 38 | $\lambda_{I_{10}, M_2}$ | 0 | 0 | 0 | 0 | 0 | 0 | 0 | 0 |

**Fig. B1.**
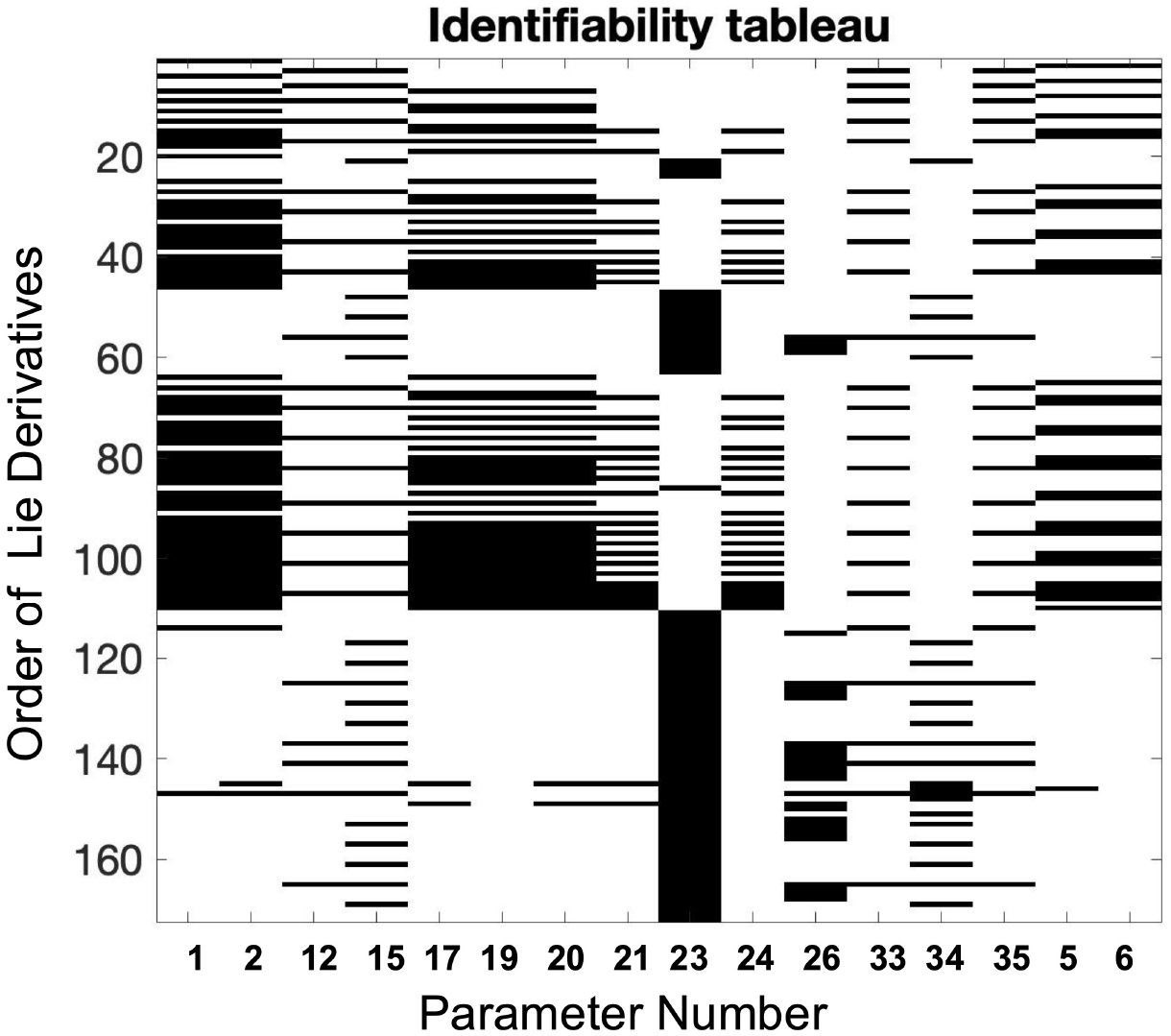
The identifiability tableau summarizes the Lie derivative used to achieve identifiability for the parameters with respect to the model outputs.

